# Magnetic nanoparticle enrichment reveals conserved nitric oxide burst responses in *Arabidopsis* and wheat

**DOI:** 10.64898/2026.11.12.765547

**Authors:** Ruixue Ji, Ning Yang, Luyao Yu, Wei Liu, Menghuan Qu, Pengcheng Wang, Guochen Qin

## Abstract

Rapid nitric oxide bursts are a conserved response to environmental stimuli, but their effects on cellular S-nitrosylation remain poorly understood. We developed a fluorinated magnetic nanoparticle method for rapid, efficient profiling of S-nitrosylation. Using *Arabidopsis* and wheat, we treated plant tissues with S-nitrosoglutathione (GSNO) to simulate a rapid nitric oxide surge. The method was then employed to capture S-nitrosylation dynamics within 24 h after treatment. Our analysis revealed sequence-dependent preferences for cysteines near basic or hydrophobic residues. Proteins showing rapid S-nitrosylation increases were predominantly associated with oxidative stress responses and energy metabolism, consistent with our metabolomic data. Functional studies using wheat *TaGR1* mutants identified glutathione reductase 1 as a key regulated target. Modification at a conserved cysteine site rapidly modulated cellular glutathione levels in both species. Here, we provide an analytical tool and a multi-omics resource for investigating S-nitrosylation dynamics.

## INTRODUCTION

Nitric oxide (NO) is a ubiquitous signaling molecule that regulates diverse physiological processes in plants and animals^1–2^. The biological effects of NO arise from its chemical reactions with molecules ranging from molecular oxygen to proteins^1,3^. One mechanism by which NO regulates protein function is S-nitrosylation, a post-translational modification. Numerous studies have shown that S-nitrosylation modulates abiotic stress responses, phytohormone signaling, immune responses, and plant growth and development by altering target protein stability, activity, subcellular localization, conformation, or protein-protein interactions^4–5^. Previous studies have indicated that the extent of protein S-nitrosylation depends on NO levels *in vivo*^6–7^.

Under stress or pathogen attack, plants rapidly increase NO levels to adapt to environmental challenges^8–10^. Nevertheless, the effects of this rapid NO burst on cellular S-nitrosylation remain poorly characterized. Few studies have examined the dynamics of protein S-nitrosylation triggered by rapid fluctuations in NO concentration.

Various proteomic approaches have been developed to profile S-nitrosylation. Among these, fluorous tagging coupled with mass spectrometry and fluorous solid-phase extraction has been widely used for S-nitrosylation profiling in animal and plant samples^11–13^. Fluorous tags are hydrophobic and can improve ion fragmentation efficiency during mass spectrometry, enabling the detection of trace amounts of S-nitrosylated proteins. However, existing fluorous solid-phase extraction materials for enriching fluorous-tagged peptides, including nanographite fluoride (nGF) and fluorous NuTips, require prolonged centrifugation. These time-consuming procedures introduce considerable technical errors, making them unsuitable for monitoring dynamic S-nitrosylation changes. Here, we exploit the speed and convenience of magnetic bead-based enrichment to develop a fluorine-functionalized magnetic nanoparticle (F-MNPs) method for high-quality profiling of S-nitrosylation dynamics. Using *Arabidopsis* and wheat as models, we treat plant materials with the NO donor S-nitrosoglutathione (GSNO) to mimic rapid NO bursts. We then use the F-MNPs method to profile S-nitrosylation changes over 24 h following treatment. Further assays identify glutathione reductase 1 (GR1) as a key target of rapid NO elevation. NO mediates S-nitrosylation at a conserved GR1 site, rapidly modulating cellular glutathione (GSH) levels. Moreover, NO enhances GR1 activity through this modification. *In vitro* assays identify C73 as a putative regulatory cysteine residue conserved across plant species, including *Arabidopsis* and wheat.

## RESULTS

### Magnetic enrichment for dynamics S-nitrosoproteome profiling

In previous studies of the fluorous affinity tag-switch (FAT-switch) method for S-nitrosylation profiling, nanographite fluoride (nGF) served as the affinity matrix for enriching FAT-tagged peptides and identifying S-nitrosylated proteins^11–12^. However, the centrifugation-based nGF procedure prolongs processing and introduces considerable operator-dependent variability. To address these limitations, we designed and synthesized Fe_3_O_4_@SiO_2_@TFS magnetic nanoparticles for rapid, efficient enrichment of FAT-tagged peptides. We termed this enrichment workflow F-MNPs (Supplementary Fig. 1). The design eliminates repeated centrifugation steps, reduces sample handling, and accelerates enrichment, providing a more robust and scalable platform for S-nitrosoproteomic analysis.

In this study, we synthesized fluorine-containing silica magnetic nanoparticles (Fe_3_O_4_@SiO_2_@TFS) using a one-step reaction in an NH_3.H2_O-ethanol solution, in which silica magnetic nanoparticles (Fe_3_O_4_@SiO_2_) were reacted with trimethoxy(1H,1H,2H,2H-nonafluorohexyl) silane (TFS)^14–15^ (Fig. 1a). This reaction rapidly yields Fe_3_O_4_@SiO_2_@TFS for enrichment of FAT-labeled peptides.

**Fig. 1.**
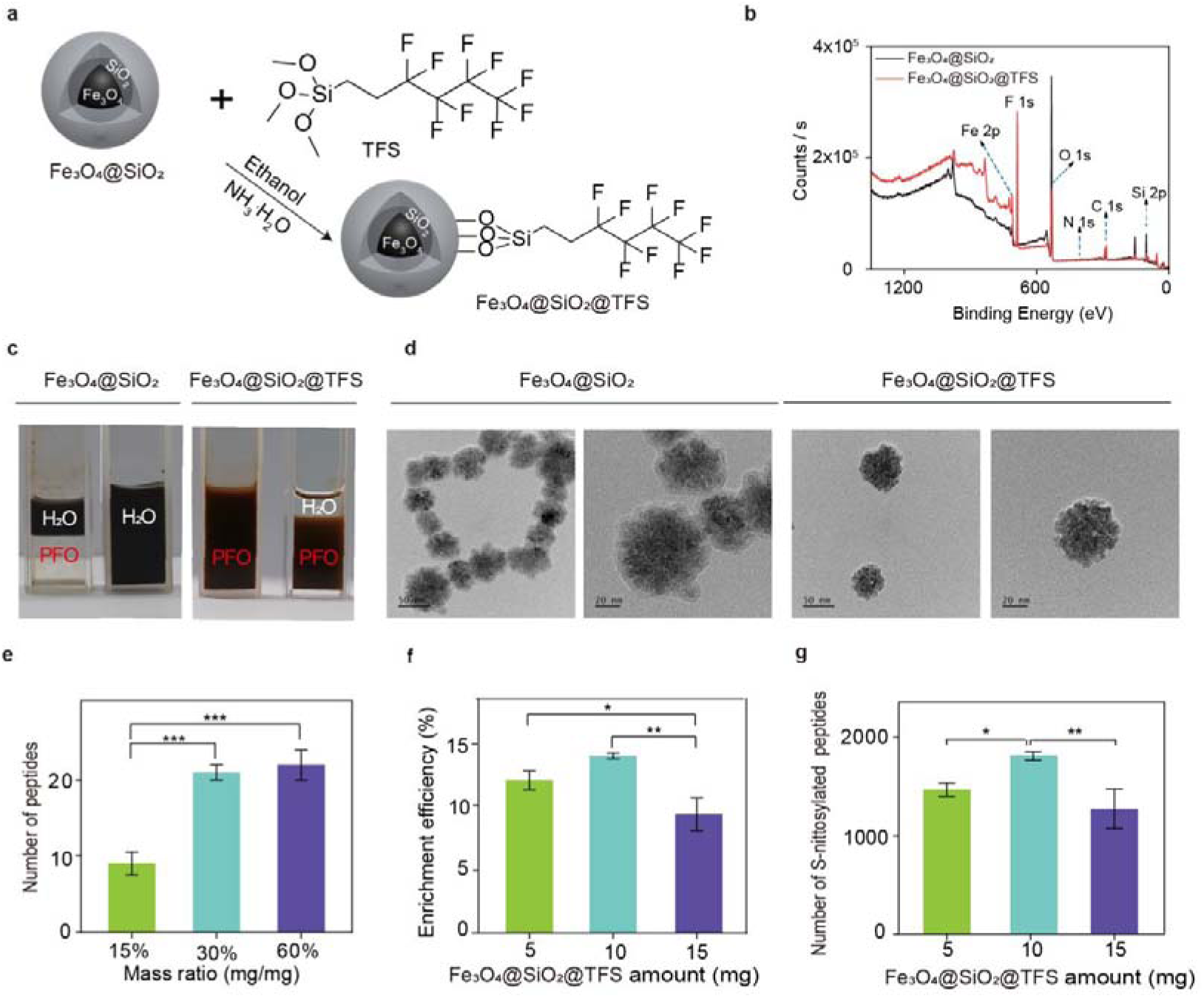
Development of the F-MNPs method. **a** Schematic illustration of fluoro-containing silica nanoparticles (Fe_3_O_4_@SiO_2_@TFS) synthesis by coating magnetic core with a layer of silica (Fe_3_O_4_@SiO_2_). **b** XPS survey spectra of Fe_3_O_4_@SiO_2_ (black line) and Fe_3_O_4_@SiO_2_@TFS (red line). **c** Photos of Fe_3_O_4_@SiO_2_ in H_2_O and H_2_O-perfluoroctane mixture solvent (left) and Fe_3_O_4_@SiO_2_@TFS in perfluoroctane and H_2_O-perfluoroctane mixture solvent (right). **d** The TEM image of Fe_3_O_4_@SiO_2_ and Fe_3_O_4_@SiO_2_@TFS. Optimization of Fe_3_O_4_@SiO_2_@TFS magnetic nanoparticles for FAT-labeled peptide enrichment. **e** Mass ratio of TFS: Fe_3_O_4_@SiO_2_ (%, w/w) on enrichment efficiency of peptides from BSA. Error bars, SD (n = 3 biological replicates). *** *p* < 0.005, one-way ANOVA. **f** The enrichment efficiency of FAT-labeled peptides from *Arabidopsis* using different amounts of Fe_3_O_4_@SiO_2_@TFS. Error bars, SD (n = 3 biological replicates). * *p* < 0.05, ** *p* < 0.01, one-way ANOVA. **g** The number of FAT-labeled peptides enriched from *Arabidopsis* seedlings by different amounts of Fe_3_O_4_@SiO_2_@TFS. Error bars, SD (n = 3 biological replicates). * *p* < 0.05, ** *p* < 0.01, one-way ANOVA.

Comparison of the X-ray Photoelectron Spectroscopy (XPS) spectra in Fig. 1b for Fe_3_O_4_@SiO_2_ (black line) and Fe_3_O_4_@SiO_2_@TFS (red line) shows a peak at 687.84 eV attributable to F 1s. The surface atomic ratios of F, C, and Si (N_F_/N_C_/N_Si_) changed from 0.48/9.13/23.08 to 33.81/24.45/5.18 (Supplementary Table S1). As shown in Fig. 1c, Fe_3_O_4_@SiO_2_ was soluble in H_2_O, but insoluble in perfluorooctane (left). As expected, Fe_3_O_4_@SiO_2_@TFS was soluble in perfluorooctane but insoluble in H_2_O (Fig. 1c). Transmission electron microscope (TEM) image show differences between Fe_3_O_4_@SiO_2_ and Fe_3_O_4_@SiO_2_@TFS (Fig. 1d). The images reveal that unmodified Fe O @SiO particles have a clear core-shell morphology, with a dark Fe O core (high electron density) and a light-grey, semitransparent amorphous silica shell (low electron density), characteristic of typical mass-thickness contrast and a monodisperse, quasi-spherical geometry. After fluorination, the particles retain their core-shell integrity and overall dispersity, while the surface becomes mildly rough, suggesting that the modification primarily affects the outer silica layer.

To optimize the amount of TFS for efficient enrichment of FAT-tagged peptides, we prepared Fe_3_O_4_@SiO_2_@TFS magnetic nanoparticles with varying TFS loadings. We then evaluated each formulation using a standard FAT-tagged peptide.

Fe_3_O_4_@SiO_2_@TFS formulations were synthesized by adding 3.75 mg, 7.5 mg, or 15 mg TFS to 25 mg Fe_3_O_4_@SiO_2_, respectively. These formulations were designated 15%-F-MNPs, 30%-F-MNPs, and 60%-F-MNPs. As shown in Fig. 1e, we used TFIA (N-(4,4,5,5,6,6,7,7,8,8,9,9,9-trideca-fluorononyl) iodoacetamide) as the FAT tag to label free thiols (–SH) on bovine serum albumin (BSA) using the FAT-switch method. The FAT-labeled peptides were enriched with Fe_3_O_4_@SiO_2_@TFS and analyzed by 90 min liquid chromatography-tandem mass spectrometry (LC-MS/MS). Whereas 15%-F-MNPs and 30%-F-MNPs enriched 9.33 ± 1.53 and 21.00 ± 1.00 FAT-labeled peptides, respectively, 60%-F-MNPs showed significantly greater enrichment sensitivity, enriching 22.00 ± 2.00 FAT-labeled peptides in total from three biological replicates. Among the tested formulations, 60%-F-MNPs (prepared at a TFS-to-Fe_3_O_4_@SiO_2_ mass ratio of 60%) performed best in both the number and intensity of enriched FAT-labeled peptides. We therefore selected this material for all subsequent S-nitrosoproteome analyses and refer to it as Fe_3_O_4_@SiO_2_@TFS hereafter.

Having established that Fe_3_O_4_@SiO_2_@TFS prepared with a 60% TFS (mass ratio) provided the most efficient enrichment of FAT-tagged peptides, we next optimized the amount of material using the same batch of *Arabidopsis* tissues. As shown in Figs. 1f and g, the number of identified S-nitrosylated peptides varied with the amount of Fe_3_O_4_@SiO_2_@TFS: 5 mg yielded 1,468.33 ± 68.25 peptides (enrichment efficiency: 12.11%), 10 mg yielded 1,809.67 ± 43.52 peptides (14.05%), and 15 mg yielded 1,272.00 ± 201.41 peptides (9.43%) (Supplementary Table S2). Among the tested amounts, 10 mg consistently yielded the most identified peptides and the highest enrichment efficiency. Using the F-MNPs method, we identified 1,809.67 ± 43.52 S-nitrosylated peptides for three replicates in this study, comparable to the 1,773.67 ± 140.00 S-nitrosylated peptides reported in the previous methods based on nGF enrichment^12^. The coefficient of variation across three replicates (CV = 2.41%) in this study was markedly lower than that of the previous dataset (CV = 7.89%), indicating better reproducibility of S-nitrosylated peptide identification in the F-MNPs method (Supplementary Table S3). Finally, the F-MNPs protocol offers a practical alternative for large-scale or time-sensitive studies that prioritize rapid processing, minimal hands-on time, and compatibility with automation.

### S-nitrosylation dynamics during nitric oxide burst

To monitor S-nitrosoproteome dynamics during an NO burst, we treated 9-day-old *Arabidopsis* seedlings with exogenous GSNO at a final concentration of 150 μM in liquid medium. GSNO is an effective NO donor with a short half-life (∼7 h)^16^, enabling rapid the robust NO release. Seedlings were harvested at 0, 0.5, 2, and 24 h after treatment, and S-nitrosylated peptides were enriched using the F-MNPs workflow for subsequent proteomic analysis.

After GSNO treatment, the number of S-nitrosylated sites in *Arabidopsis* increased rapidly at 0.5 h (supported by 2,113 peptides), decreased markedly at 2 h, and returned to the basal level by 24 h, comparable to the untreated control (0 h) (Fig. 2a; Supplementary Data 1). Principal component analysis (PCA) of the enriched S-nitrosylated sites supported these temporal dynamics: the 24 h samples clustered closely with the 0 h controls, indicating recovery to the pretreatment state, whereas the 0.5 h samples were clearly separated from the other time points, reflecting a distinct proteomic configuration at peak S-nitrosylation (Supplementary Fig. S2a).

**Fig. 2.**
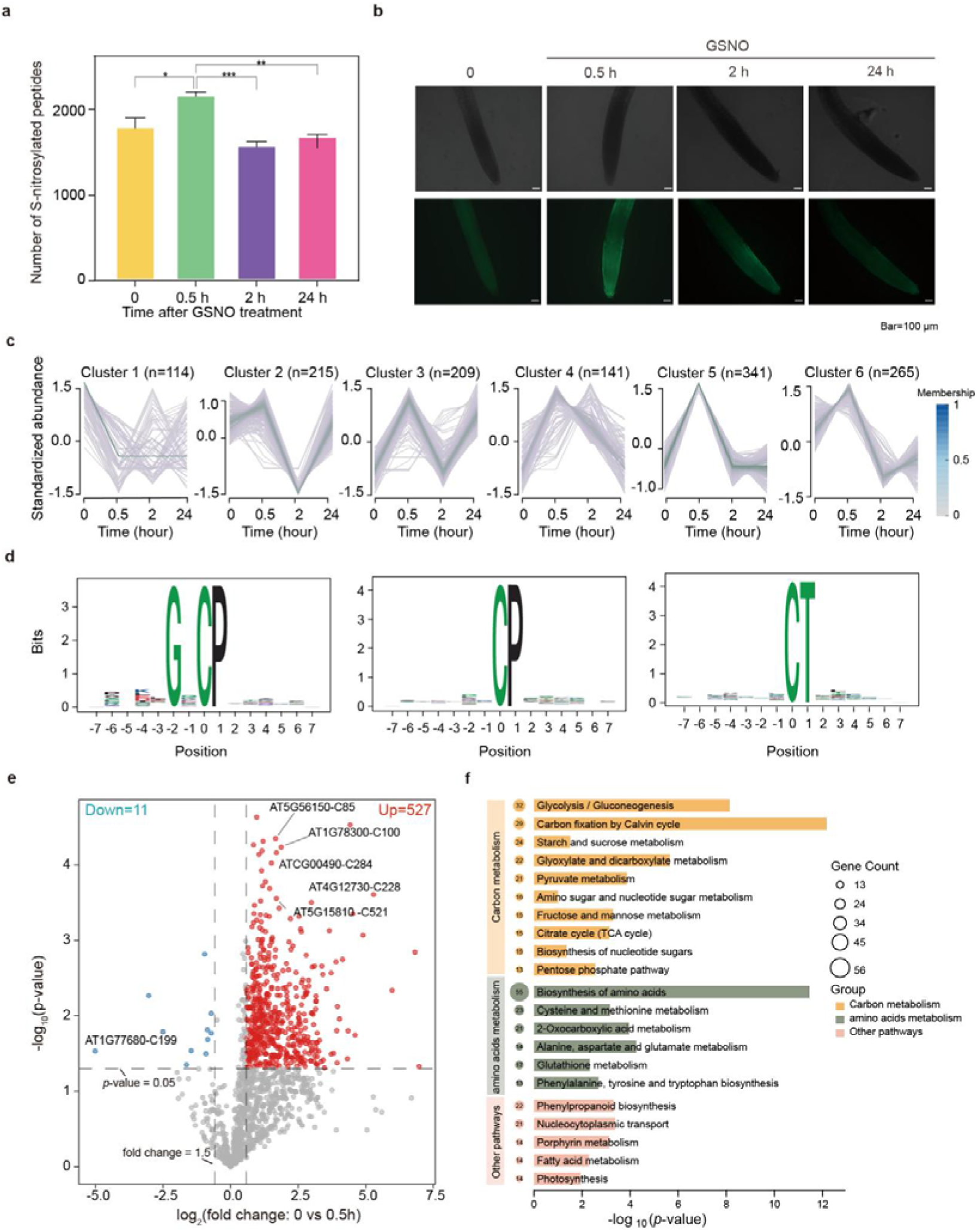
Profiling of S-nitrosylation dynamics in *Arabidopsis* by GSNO treatment. **a** The number of S-nitrosylated peptides identified in *Arabidopsis* by GSNO treatment. Error bars, SD (n = 3 biological replicates). * *p* < 0.05, ** *p* < 0.01, *** *p* < 0.005, one-way ANOVA. **b** Confocal microscopy of 9-day-old *Arabidopsis* root of GSNO treatment using DAF-FM to NO distribution. **c** Specific expression pattern in the GSNO treatment proteome dataset. Six cluster: 114 sites shows down-remainten (cluster 1). 215 sites were up-down-up (cluster 2). 209 sites were up-down-up (cluster 3). 141 sites were up-down-down (cluster 4). 341 sites were up-down-hold (cluster 5). 265 sites were up-down-up (cluster 6). **d** Motif analysis of 1171 S-nitrosylation sites which were up-regulated at 0.5 h-post GSNO treatment. Over representation motif of Cys-SNO sequence windows (-7 to +7 residues) constructed by rmotifx. **e** Volcano plot of S-nitrosylation site ifentify in 0.5 h post-GSNO treatment and control conditions. Red dots (n = 527) represent S-nitrosylated sites that were up-regulated at 0.5 h post-GSNO treatment, blue dots (n = 11) represent S-nitrosylated sites that were down-regulated at 0.5 h post-GSNO treatment (|fold change| > 1.5, *p* < 0.05, two-tailed unpaired *t* test). **f** KEGG pathway analysis of upregulated S-nitrosylated proteins in *Arabidopsis* after 0.5 h of GSNO treatment.

These findings indicate that GSNO treatment transiently increases protein S-nitrosylation in *Arabidopsis*. Consistent with this pattern, confocal laser scanning microscopy (CLSM) using the NO-sensitive fluorescent probe DAF-FM revealed a clear increase in fluorescence at 0.5 h, followed by a decrease. By contrast, GSH treatment did not significantly enhance fluorescence in CLSM imags (Supplementary Fig. S2b).

Based on the S-nitrosoproteomic data, we conducted cluster analysis of 1,285 S-nitrosylated sites showing significant changes (adjusted *p*-value < 0.05). This analysis classified the sites into six clusters according to their abundance trends (Fig. 2c).

Cluster 1 (114 sites) showed significant downregulation at 0.5 h after GSNO treatment. Cluster 2 (215 sites), Cluster 3 (209 sites), Cluster 4 (141 sites), Cluster 5 (341 sites) and Cluster 6 (265 sites) were significantly upregulated at 0.5 h. These findings are consistent with the PCA results. Given that most clusters exhibited higher signal intensity after 0.5 h of GSNO treatment, we focused our functional analysis on clusters 2-6. We performed motif analysis on 1,171 S-nitrosylated sites to characterize the site-specific frequencies of amino-acid residues surrounding S-nitrosylated cysteines (Fig. 2d, Supplementary Fig. S3). For motif analysis, protein sequences spanning seven amino acids upstream and downstream of each S-nitrosylation site were extracted to generate 15 amino-acid windows centered on the modified residue. Enriched sequence motifs were identified using rmotifx^17^. Motif analysis of SNO sites in clusters 2-6 identified 13 enriched motifs, with significant enrichment of motifs containing hydrophobic (valine, V; proline, P; alanine, A) or basic residues (lysine, K; arginine, R). The enriched motifs included G-x-C-P, C-P, C-T, C-G-x-x-x-P, K-x-C-A, C-x-x-x-x-R, V-x-x-x-x-x-x-C, C-x-x-x-x-x-A, C-x-x-A, Q-x-C, C-x-G-x-R, C-G, and C-x-x-x-x-x-K. This pattern reflects the greater susceptibility to SNO modification of cysteine residues with nearby basic motifs, greater sulfur-atom exposure, surrounding hydrophobicity, and a low pKa^2,12,18–20^.

We next performed a pairwise comparison of S-nitrosylated sites between *Arabidopsis* at 0.5 h after GSNO treatment and untreated control (0 h). In total, 527 S-nitrosylated sites were significantly upregulated in *Col_0* samples at this time point, whereas only 11 were downregulated (|fold change| > 1.5, *p* < 0.05, two-tailed unpaired *t-test*) (Fig. 2e). These findings suggest that NO signaling following GSNO treatment rapidly induces S-nitrosylation.

To delineate the earliest proteomic events triggered by GSNO before secondary responses, we performed Gene Ontology (GO) enrichment analysis of *Arabidopsis* S-nitrosylated proteins differentially expressed at 0.5 h (Supplementary Fig. S4). Early biological processes showed marked enrichment in photosynthesis, hydrogen peroxide catabolism, and responses to oxidative stress. Concurrently, cellular component annotations were dominated by chloroplast-related terms (stroma, envelope, plastid), cytosol, and cytoplasm. Metal ion binding, ATP binding, and protein binding were the most prominently enriched molecular function categories.

This functional profile indicates that the 0.5 h GSNO proteome rapidly reconfigures energy metabolism and antioxidant defense, particularly organellar (especially chloroplastic) and redox-regulatory processes. Collectively, these data reveal an early metabolic alert phase in which GSNO-mediated NO signaling fine-tunes photosynthetic carbon assimilation and oxidative stress management, laying the foundation for more complex, organ-specific proteomic adaptations at later time points. This temporal resolution refines our understanding of NO as a rapid, multi-compartmental signal that prioritizes cellular redox and energy homeostasis during early exposure.

We next identified enriched Kyoto Encyclopedia of Genes and Genomes (KEGG) pathways among proteins with increased S-nitrosylation in *Arabidopsis* at 0.5 h after GSNO treatment (Fig. 2f). The significantly enriched pathways fell into two major KEGG categories: (i) carbohydrate metabolism, encompassing carbon metabolism, glycolysis/gluconeogenesis, the Calvin cycle (carbon fixation), pyruvate metabolism, and glyoxylate and dicarboxylate metabolism; and (ii) amino-acid metabolism, including amino-acid biosynthesis and cysteine/methionine metabolism, together with the closely associated biosynthesis of other secondary metabolites.

### Metabolic remodeling during nitric oxide burst

Dynamic S-nitrosylation proteomics revealed that an NO burst rapidly increases S-nitrosylation of proteins in multiple vital metabolic pathways. Whether these modifications alter metabolite abundance in the corresponding pathways warrants further investigation. Given previous reports that NO plays a critical role in the regulation of plant metabolism^21–24^, we performed untargeted metabolomic analysis of *Arabidopsis* seedlings at 0, 6, and 24 h after GSNO treatment. Classification of the 391 detected metabolites showed relatively high proportions of amino acids and their derivatives, organic acids, and carbohydrates (Fig. 3a; Supplementary Data 2). PCA revealed treatment-duration-dependent fluctuations in metabolite profiles (Fig. 3b).

**Fig. 3.**
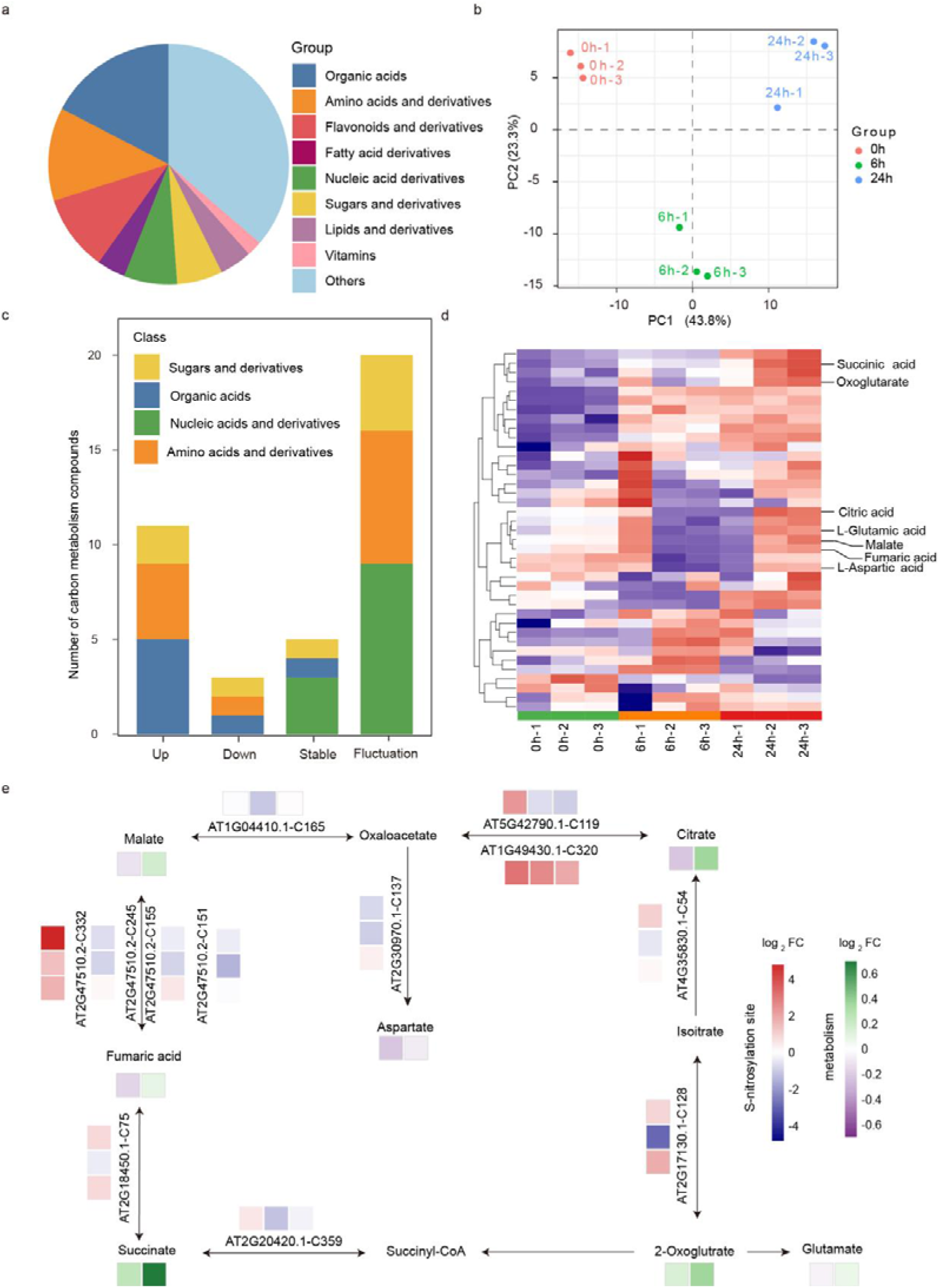
Metabolomic changes in *Arabidopsis* seedlings after GSNO treatment. **a** Untargeted metabolomic analysis of GSNO-treated *Arabidopsis* seedlings: statistics of detected metabolites and their classification. **b** Principal component analysis of metabolites from the *Arabidopsis* under GSNO treatment for 0 h, 6 h, and 24 h. **c** Statistics of carbon metabolites in different dynamic change patterns from the *Arabidopsis* under GSNO treatment for 0 h, 6 h, and 24 h. **d** Heatmap of compounds in carbon metabolism from the *Arabidopsis* under GSNO treatment for 0 h, 6 h, and 24 h. **e** Metabolic network analysis of TCA cycle showing GSNO-induced changes in metabolites and proteins site intensities. Metabolite abundances and site intensities were expressed as log_2_-fold changes relatives to the 0 h baseline, calculated from mean signal intensities. Rows represents metabolites or protein sites; from left to right, the GSNO treatment time points (6 h, and 24 h for metabolites; 0.5, 2 and 24 h for protein sites). Green and red indicated increased abundance relative to 0 h, and pruple and blue indicates decreases.

Analysis of metabolite dynamics showed that most compounds fluctuated over time. Specifically, 81 metabolites (20.72%) increased persistently, 33 (8.44%) decreased persistently, 202 (51.66%) fluctuated, and 75 (19.18%) remained stable. These findings indicate GSNO-induced metabolic remodeling in *Arabidopsis* (Supplementary Fig. S5a).

To further examine how NO-burst-induced S-nitrosylation regulates metabolic activities and reshapes metabolite abundance in plants, we focused on carbon metabolism. Untargeted metabolomic analysis identified 39 metabolites involved in carbon metabolism, of which 87.18% changed in abundance within 24 h of GSNO treatment. This finding strongly supports our S-nitrosylation proteomic data, which revealed that 73 proteins in carbon metabolism pathways were S-nitrosylated, including several key regulators (Supplementary Table S4).

The tricarboxylic acid (TCA) cycle is the central hub of carbon metabolism^25^. In Fig. 3e, nine enzymes participating in the TCA cycle underwent S-nitrosylation, five metabolites were detected by mass spectrometry, and their levels also changed within 24 h after GSNO treatment. Because the biosynthetic precursors of glutamate and aspartate derive from the TCA cycle, changes in intermediate abundance also alter the biosynthesis of these amino acids. Collectively, our results demonstrate that NO bursts alter protein S-nitrosylation in multiple basal metabolic pathways, subsequently leading to metabolic reprogramming in plant cells.

### Conserved nitric oxide burst response in wheat

Based on these findings in *Arabidopsis*, we extended our investigation to wheat (*Triticum aestivum L.* cv. Fielder), an important cereal crop. To examine whether GSNO-mediated protein S-nitrosylation is conserved in monocots, we applied the same experimental regime and performed proteomic profiling of GSNO-treated wheat seedlings (Supplementary Data 3). Using the proteomic data, we conducted cluster analysis of 1,421 sites. This analysis classified the sites into six clusters according to their abundance trends (Fig. 4a). Cluster 1 (256 sites), Cluster 2 (129 sites), Cluster 4 (284 sites), and Cluster 5 (273 sites) were upregulated at 1 h, whereas Cluster 3 (332 sites) and Cluster 6 (147 sites) remained at baseline. Cluster analysis of S-nitrosylated sites showed that many sites increased rapidly in abundance at 1 h after GSNO treatment in wheat seedlings. This finding was supported by the volcano plot, which showed 1,247 significantly upregulated S-nitrosylated sites and only 15 sites downregulated at 1 h after treatment (Supplementary Fig. S6a). Together, these findings indicate that GSNO can rapidly induce S-nitrosylation, which may underlie early metabolic and signaling responses in wheat seedlings.

**Fig. 4.**
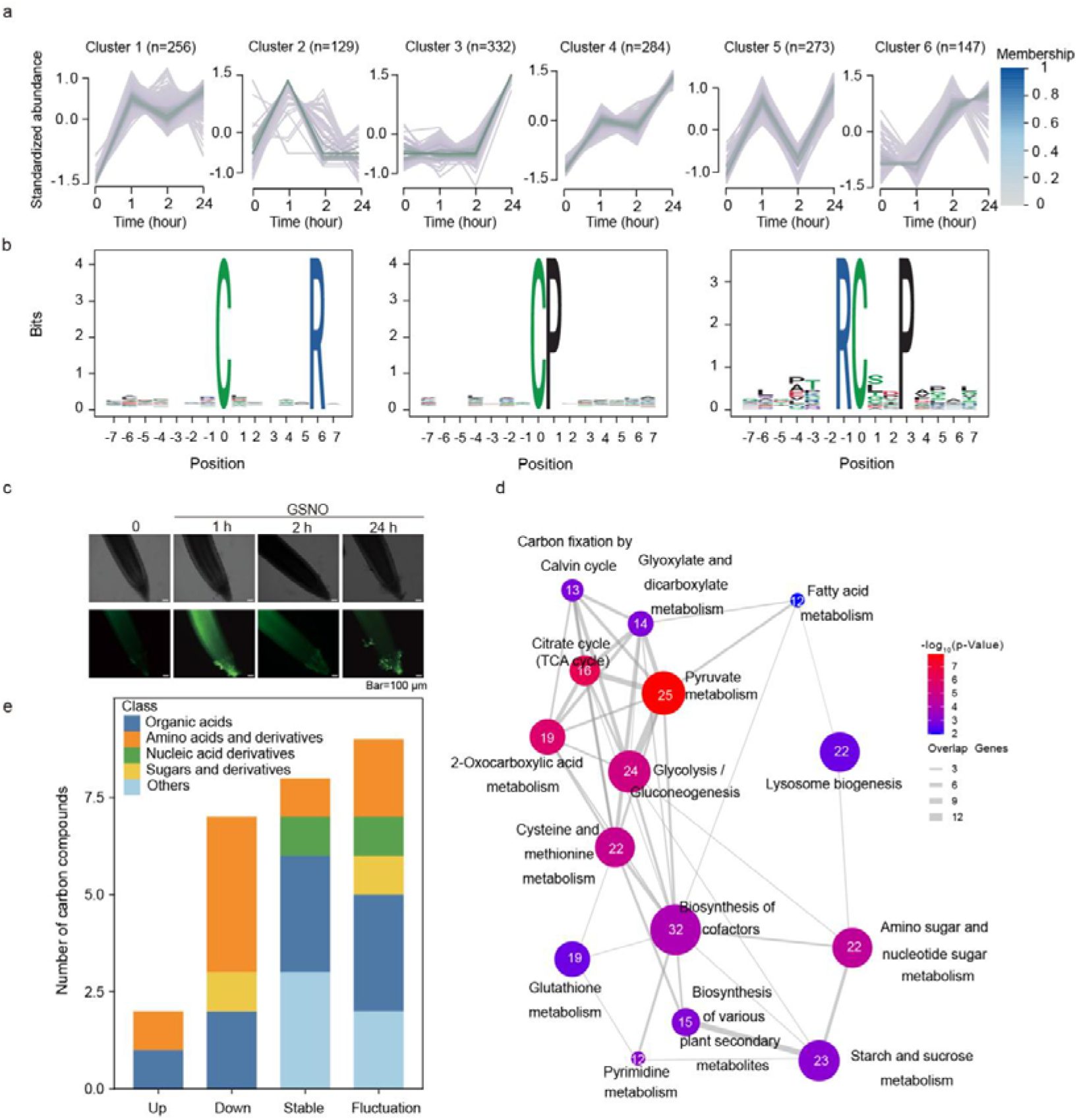
Profiling of S-nitrosylation dynamics in wild-type wheat following GSNO treatment. **a** Specific expression pattern in the GSNO treatment proteome dataset. Six cluster trends: Cluster 1 (256 sites) is up-down-up. Cluster 2 (129 sites) is up-down-hold. Cluster 3 (332 sites) is hold-hold-up. Cluster 4 (284 sites) is up-hold-up. Cluster 5 (273 sites) is up-down-up. Cluster 6 (147 sites) is hold-up-up. **b** Motif analysis of 942 S-nitrosylation sites enriched in Fielder. Over representation motif of Cys-SNO sequence windows (-7 to +7 residues) constructed by rmotifx. **c** Confocal microscopy of 5-day-old Fielder root of GSNO treatment using DAF-FM to NO distribution. **d** KEGG pathway analysis of upregulated S-nitrosylated proteins in Fielder after 1 h of GSNO treatment. **e** Stacked bar chart showing the classification and change trends of compounds assigned to the carbon metabolism pathway.

Trend analysis revealed that the majority of clusters exhibited increased signal intensity upon GSNO treatment for 1 h. therefore, we performed functional analysis on clusters 1, 2, 4, and 5 in Fig. 4a. To further characterize the S-nitrosylation sites, we used rmotifx to compare position-specific amino-acid frequencies surrounding S-nitrosylated cysteines across 942 sites (Fig. 4b, Supplementary Fig. S6b). The analysis showed significant enrichment of motifs containing hydrophobic residues (valine, V; proline, P; alanine, A) or basic residues (lysine, K; arginine, R). The enriched motifs included C-x-x-x-x-x-R, C-P, R-C-x-x-P, P-x-C, C-x-x-x-x-R, K-x-x-x-x-x-x-C, C-x-G, C-x-x-x-x-K, and C-x-x-P. Analysis of SNO sites across these groups revealed higher frequencies of hydrophobic residues (proline, P) and basic residues (lysine, K; arginine, R). This motif signature closely matches the patterns observed in *Arabidopsis* following NO donor treatment.

Compared with GSH treatment, GSNO treatment produced a more pronounced change in confocal fluorescence (Fig. 4c, Supplementary Fig. S6c). Confocal microscopy further supported this early response, showing enhanced fluorescence at 1 h and confirming rapid accumulation of S-nitrosylated sites *in vivo*. Fluorescence intensity was significantly higher than in the control at 1 h, then progressively decreased from 2 h onward. Consistent with our proteomic analyses, GSNO treatment rapidly elicited protein S-nitrosylation. GO analyses also revealed significant enrichment of biological processes related to oxidative stress response and carbohydrate metabolism (Supplementary Fig. S6d). KEGG pathway analysis further indicated that these GSNO-responsive proteins were primarily involved in carbohydrate and energy metabolism (Fig. 4d). Carbon metabolism, pyruvate metabolism, glycolysis/gluconeogenesis, and glutathione metabolism were notably overrepresented among the identified proteins. Untargeted metabolomic profiling of GSNO-treated wheat detected 396 compounds. As in *Arabidopsis*, most compounds were amino acids, organic acids, and sugars, and PCA revealed comparable patterns (Supplementary Fig. S7a and b; Supplementary Data 4). Building on the *Arabidopsis* findings, we specifically examined carbon metabolic remodeling in wheat. After GSNO treatment, most carbon metabolism-related compounds fluctuated in abundance (Fig. 4e). These results suggest that exogenous GSNO reshapes plant energy and carbon flux distribution through S-nitrosylation, facilitating adaptation to environmental changes. Comparison with untargeted metabolomic data from *Arabidopsis* revealed broadly similar changes in carbon metabolism-related compounds after GSNO treatment (Supplementary Figs. S7c and d). Although these compounds showed more complex dynamics in wheat than in *Arabidopsis*, key nodal metabolites in carbon metabolism followed consistent trends.

### GR1 S-nitrosylation rapidly regulates glutathione levels

Glutathione (GSH) is a major plant antioxidant^26–27^. Following GSNO treatment, proteins involved in GSH biosynthesis and metabolism were rapidly S-nitrosylated in both *Arabidopsis* and wheat (Supplementary Table S5). Consistent with this finding, metabolomic analysis revealed a rapid increase in GSH abundance in both species after GSNO exposure (Supplementary Figs. S8a and b). We also found that GR1, a key enzyme catalyzing the reduction of oxidized glutathione (GSSG) to GSH, was S-nitrosylated in both *Arabidopsis* and wheat. *Arabidopsis* GR1 C73 and wheat GR1 C71 were both S-nitrosylated (Figs. 5a and b), and the corresponding modified peptides increased rapidly in abundance by 1.15-fold and 4.04-fold, respectively, after GSNO treatment (Supplementary Table S6). These results indicate that GR1 serves as a critical target regulated by NO.

**Fig. 5.**
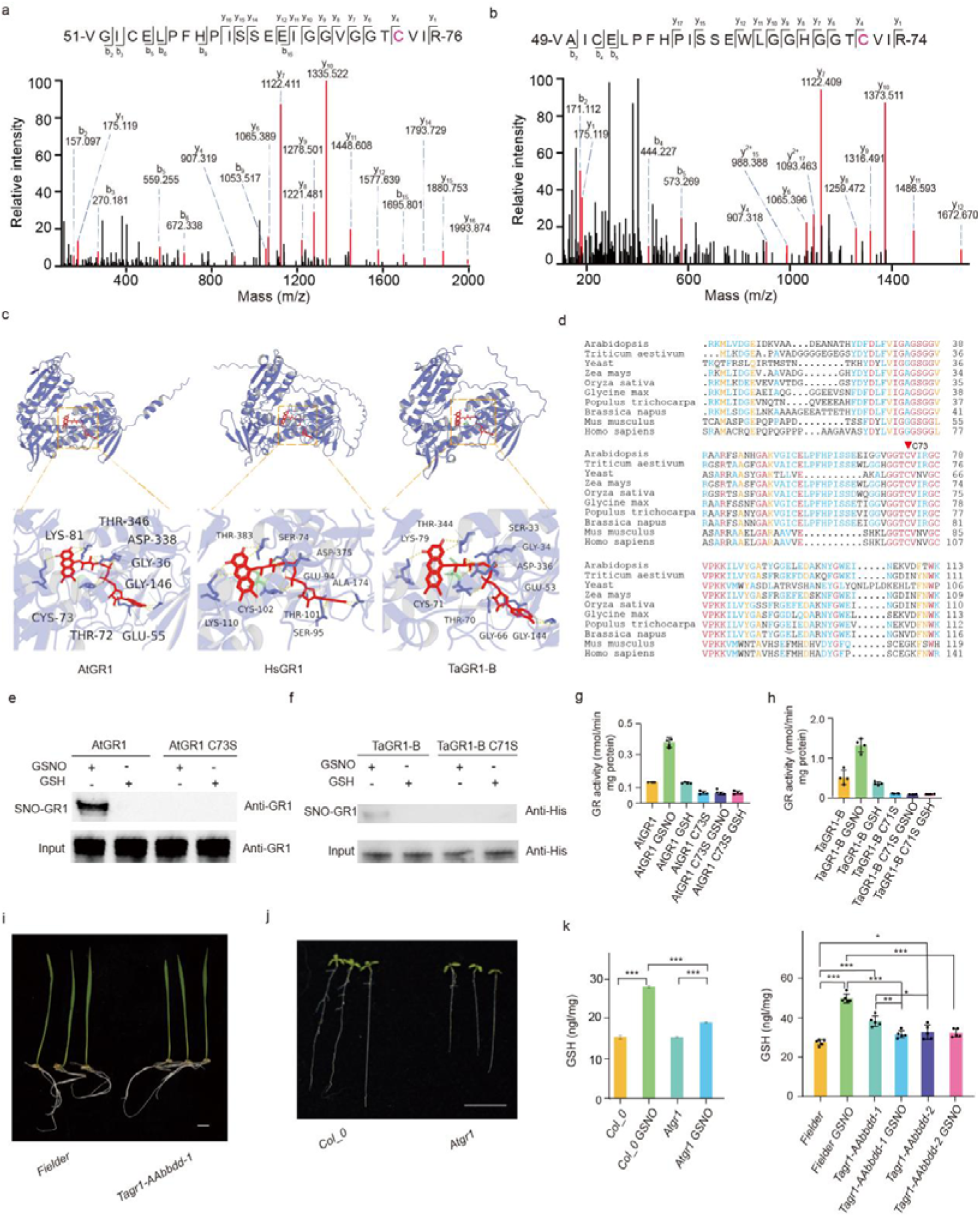
S-nitrosylation enhance GR1 activity. **a** The MS/MS spectrum of S-nitrosylated peptide containing Cys73 in AtGR1. **b** The MS/MS spectrum of S-nitrosylated peptide containing Cys71 in TaGR1-B. **c** Structural modeling of AtGR1, TaGR1-B and HsGR1. Red sticks represent the FAD molecule. The disulfides C71 in TaGR1-B, and C73 in AtGR1 are highlighted. **d** Alignment of GR1 protein sequences in different model species. The alignment was performed with MUSCLE (https://www.ebi.ac.uk/Tools/msa/muscle).The cysteine residues corresponding to active sites are highlighted by red arrows. **e** Analysis of S-nitrosylation of recombinant wild-type and Cys73Ser mutated AtGR1 without or without GSNO/GSH treatment. The S-nitrosylated recombinant AtGR1 and the loading of AtGR1 and AtGR1^C73S^ was measured by ware detected by anti-GR immunoblot. **f** Analysis of S-nitrosylation of recombinant wild-type and Cys71Ser mutated TaGR1-B with or without GSNO treatment. The S-nitrosylated recombinant TaGR1 and the loading of His-TaGR1 and His-TaGR1^C71S^ was measured by ware detected by anti-His immunoblot. **g** Analysis of AtGR1 and AtGR1^C73S^ activities with or without GSNO treatment by Glutathione Reductase Assay Kit with DTNB [5,5-Dithio-bis-(2-nitrobenzoic acid)] colorimetry. **h** Analysis of TaGR1 and TaGR1^C71S^ activities without or with GSNO treatment by Glutathione Reductase Assay Kit with DTNB [5,5-Dithio-bis-(2-nitrobenzoic acid)] colorimetry. **i** Phenotype of *Fielder* (wild type) and *Tagr1-AAbbdd-1* (bar = 1 cm). Photograph were taken after growing at 22 °C for 4 days. **j** Phenotype of *Col_0* (wild type) and *Atgr1* (bar = 1 cm). Photograph were taken after growing at 22 °C for 9 days. **k** GSH content of *Col_0* (wild type) and *Atgr1* plants with or without GSNO treatment. Error bars, SD (n = 3). *** *p* < 0.005, one-way ANOVA. **l** GSH content of *Fielder* (wild type), *Tagr1-AAbbdd-1* and *Tagr1-AAbbdd-2* plants with or without GSNO treatment. Error bars, SD (n = 5). \**p* < 0.05, \*\**p* < 0.01, *\*\*\* p* < 0.005, one-way ANOVA.

Structural analysis of GR1 revealed that C73 in AtGR1, C71 in TaGR1, and C102 in HsGR1 lie within the flavin adenine dinucleotide (FAD)-binding pocket (Fig. 5c), a potentially important region for activity regulation. Evolutionary analysis further showed that the regions encompassing C73 in *Arabidopsis*, C71 in wheat, and C102 in humans are highly conserved among the three species (Fig. 5d). We purified recombinant *Arabidopsis* and wheat GR1 proteins *in vitro* and generated corresponding cysteine-to-serine (Cys-to-Ser) mutant fusion proteins at C73 and C71, respectively. *In vitro* assays confirmed C73 and C71 as the primary GR1 S-nitrosylation site (Figs. 5e and f). Enzyme assays showed that GSNO treatment rapidly enhanced GR1 activity, whereas mutation of C73 and C71 abolished this activation (Figs. 5g and h).

We next generated a wheat GR1 double mutant, AAbbdd, using CRISPR/Cas9 gene editing (Supplementary Figs. S9a-c; Fig. 5i). This mutant and the *Arabidopsis gr1* mutant (Supplementary Fig. S9d; Fig. 5j) were used in GSNO treatment assays.

Under normal conditions, the *Arabidopsis gr1* mutant accumulated more GSH than the wild type, consistent with previous reports^26^. After GSNO treatment, GSH levels increased in wild type *Arabidopsis* but declined in the *gr1* mutant (Fig. 5k). In wheat, basal GSH concentrations did not differ significantly between the *GR1* AAbbdd mutant and the wild type. However, GSNO treatment markedly increased GSH in wild-type wheat, whereas this response was absent in the mutant (Fig. 5l).

Collectively, these data demonstrate that GR1 is a major S-nitrosylation target during rapid NO bursts and a key driver of increased cellular GSH.

## DISCUSSION

NO bursts are a widespread intrinsic molecular signaling events involved in adaptation to environmental fluctuations. Rapid NO elevation in response to diverse external stimuli occurs in plants, animals, and humans^7–10^. Nevertheless, how rapid NO accumulation drives cellular S-nitrosylation dynamics remains poorly understood, including which protein sites are initially modified and which signaling pathways are activated first. These unresolved questions motivated this study. We used GSNO, the most common physiological NO reservoir in nature, as an exogenous donor to mimic NO bursts. Integrating proteomic and metabolomic approaches, we characterized the dynamic effects of rapid NO bursts on S-nitrosylation and cellular metabolites.

Notably, our results showed that, under nonenzymatic conditions, S-nitrosylation preferentially targets cysteines adjacent to hydrophobic and basic residues. The affected pathways predominantly involve amino-acid biosynthesis and carbon metabolism. This finding advances the sequence-level identification of rapid NO-regulated sites and offers insights for future rapid screening of NO targets using artificial intelligence.

GSH metabolism and biosynthesis form a core antioxidant metabolic pathway in living organisms. In plants, this pathway is required for responses to diverse environmental stresses. GR proteins catalyze the reduction of oxidized GSSG to GSH and are primary regulators of rapid GSH production *in vivo*^28–30^. Numerous studies have linked GR activity to plant stress responses and shown that NO regulates this activity. However, the key regulatory cysteine sites in GR proteins have not yet been identified³¹^-^³³. Our analyses in *Arabidopsis* and wheat demonstrate that C73 in the C-terminal region of GR is critical for regulating GR activity. This residue lies at the FAD cofactor-binding interface and is an immediate target for rapid NO modification. This finding provides insight into the mechanism regulating GR activity. Our proteomic data also reveal additional S-nitrosylation changes in other GSH metabolism-associated proteins. Further investigation of their regulatory relationships with NO will help elucidate the broader NO signaling network.

In this work, we combined fluorous solid-phase extraction with rapid magnetic bead enrichment to develop the F-MNPs method for time-resolved profiling of S-nitrosylation over 24 h under NO burst conditions. By integrating metabolomics, we further characterized how rapid NO elevation reshapes plant metabolism and modulates adaptive responses to environmental cues. We investigated NO bursts in both the model plant *Arabidopsis thaliana* and wheat, an agriculturally important crop. Using CRISPR/Cas9 gene editing, we generated *GR1* mutants to dissect how NO bursts rapidly alter GR1 activity and thereby modulate cellular GSH levels. Because GSH is a vital antioxidant for plant stress tolerance, these findings have potential applications in crop breeding.

## METHODS

### Plant materials and growth conditions

The *Arabidopsis* wild type *Col_0* and atgr1 mutant (SALK_104319C) seeds were sterilized with 75% ethanol, and then seeds were germinated in 100 mL of 1/2 MS liquid medium containing 0.5% (w/v) sucrose after a 4-day 4 °C treatment. Seedlings were grown at 22 °C with 16 h light / 8 h dark for 9 d before being harvested for S-nitrosoproteomic analysis. Then, 15 μL of 1 M GSNO into 100 mL 1/2 MS liquid medium, and seedlings were grown at 22 °C with 0, 0.5, 2 and 24 h after treatment.

The wheat (*Triticum aestivum*) Fielder seeds were sterilized with 75% ethanol solution, and then seeds were germinated in 10 mL H_2_O after 4-day 4 °C treatment. Seedlings were grown at 22 °C with 16 h light for 5 d before being harvested for S-nitrosoproteomics assay. Then, 15 μL of 0.1 M GSNO was added to 10 mL of H_2_O, seedling were harvested at 0, 1, 2 and 24 h after treatment.

### Preparation of Fe_3_O_4_@SiO_2_@TFS nanoparticles

The 25 mg pure **Fe_3_O_4_@SiO_2_ nanoparticles** dissolved in 5 mL of ethanol solution in a 50 mL tube. TFS (Triethoxy (1H,1H,2H,2H-nonafluorohexyl) silane) and 5 mL of ammonium hydroxide were added to the tube, the mixture was reacted at room temperature for 12 h. The samples were washed with methanol, the methanol was removed by magnetic adsorption with a magnetic rack, and after repeating the wash several times, the residue was dried under N_2_. Finally, the purified **Fe_3_O_4_@SiO_2_@TFS** was stored at 4 °C.

### *In vivo* S-nitrosylated protein enrichment

For *Arabidopsis,* 0.5 g seedling samples were homogenized in 1.5 mL of ice-cold HEN buffer (250 mM HEPES, pH 7.7, 1 mM EDTA, 0.1 mM neocuproine) containing 0.5% Triton X-100, 0.1 mM PMSF and protease inhibitor cocktail for 30 min with shaking. The protein extracts were centrifuged at 20,000 × g for 15 min at 4 °C, and the supernatant was collected. The proteins were diluted to 1 mg/mL with HEN buffer and were incubated with 3 volumes of blocking buffer (250 mM HEPES, pH 7.7, 1 mM EDTA, 0.1 mM neocuproine, 5% SDS, 270 mM IAM) at 37 °C for 120 min. The samples were precipitated by incubating with 2 volumes of acetone at-20 °C for 30 min. The pellet was washed three times with 70% acetone and dissolved in 0.5 mL of HEN buffer with 1% SDS (HENS buffer). The concentration of proteins was detected by BCA assay, and 12.5 μL of 200 mM sodium ascorbate and 400 μg of TFIA were added to samples which contained 1 mg of protein in 0.4 mL of HENS buffer. After labeling for 2 h at room temperature with shaking and centrifuging at 200 × g, the supernatant was collected; extra TFIA was removed with beads (Sera-Mag carboxylate, Cytiva). Proteins were digested using the SP3 method (10 mM DTT treatment for 90 min and 55 mM IAM treatment for 25 min, trypsin 1:50 (w/w), digestion overnight). The samples were dried in a centrifugal vacuum concentrator (Labconco, USA).

For wheat (*Triticum aestivum*) Fielder root samples, the same experimental protocol was used without modification. Specifically, 0.5 g of root tissues was subjected to the same homogenization, blocking, acetone precipitation, TFIA labeling, and SP3-based digestion steps, using the identical buffer compositions, incubation temperatures, and centrifugation conditions as described for the *Arabidopsis* seedlings. Thus, all parameters-including protein concentration adjustment, ascorbate and TFIA amounts, and protease inhibitor cocktails-were kept strictly consistent between the two plant materials.

### BSA cysteine peptides enrichment

TCEP and 400 μg of TFIA were added to samples that contained 1 mg BSA proteins in 0.4 mL of HENS buffer. After labeling for 2 h at room temperature with shaking; then the samples were centrifugured at 200 × g, the supernatant was collected; excess TFIA was removed by filter. The sample was washed twice by pipetting with 50 mM ammonium bicarbonate solution. Add trypsin dissolved in 50 mM ammonium bicarbonate solution and digest overnight at 37 °C. Upon completion of digestion, acidify with 10% FA to terminate the reaction. The samples were dried in a centrifugal vacuum concentrator (Labconco, USA).

### The Fe_3_O_4_@SiO_2_@TFS was used to enrich FAT-tagged peptides

First, 10 mg **Fe_3_O_4_@SiO_2_@TFS** was dissolved 200 μL methanol and mixed with FAT-tagged peptides which dissolved by 200 μL of 25 mM ABC. Vortexed to mix thoroughly, and let stand on the magnetic stirrer for another 5 min, then remove the solution using a magnetic rack. The tube was washed sequentially with 200 μL of 25 mM ABC and 200 μL of 60% methanol/40% 25 mM ABC. Finally, 200 μL methanol was added to elute the FAT-tagged peptides. The eluted samples were dried in a centrifugal vacuum concentrator (Labconco, USA) and redissolved in 6 μL of 0.2% FA (containing 5% ACN) solution for LC-MS/MS analysis.

### LC-MS/MS assay

Peptide samples were dissolved in 6 μL 0.1% formic acid and 5 μL was injected into the Orbitrap Eclipse Tribrid mass spectrometer (Thermo Fisher Scientific) coupled with Vanquish neo (Thermo Fisher Scientific) and FAIMS Pro Duo (Thermo Fisher Scientific). Peptides were separated on a 20 cm in-house analytical column packed with the reverse phase material ReproSil-Pur C18-AQ (1.9 μm resin; Dr. Maisch, GmbH, Tübingen, Baden-Württemberg, Germany) with a 90 min gradient from 5% to 90% (from 5% to 23% for 7 min, from 23% to 70% for 49 min, from 70% to 80% for 14 min, from 80% to 90% for 14 min, and 90% for 9 min) of 80% (v/v) acetonitrile and 0.1% (v/v) formic acid. Full MS scans were acquired with a resolution of 60,000 in the Orbitrap. The high energy collision induced dissociation (HCD) fragment ion spectra were acquired in the Orbitrap with a resolution of 15,000 at m/z 200. The following conditions were used: spray voltage of 2.2 kV, scan range of 350-1,500 m/z, ion transfer tube temperature of 320 °C, radio frequency (RF) lens of 40%, threshold of minimum signal inten sity 5.0e^4^, dynamic exclusion durations of 22 s, HCD collision energy of 35%. FAIMS mode was set at standard resolution with 45 and 65 V.

### Mass spectrometry data analysis

MS/MS data were searched against the *Arabidopsis* Information Resource (TAIR 10) protein sequences database using Proteome Discoverer 2.4 (Thermo Fisher Scientific), and the wheat proteome sequences database (IWGSC RefSeq v2.1) using Proteome Discoverer 2.4 (Thermo Fisher Scientific). The parameters were restricted to peptides digested by trypsin with a maximum of two missed cleavages allowed. Cysteine carbamidomethylation, oxidation of methionine and TFIA (C_11_F_13_H_9_NO, 418.047656 Da) of cysteine were set as variable modifications. The minimum peptide length was set at six amino acids. Mass tolerances were set up to 10 ppm for MS ions and 0.1 Da for MS/MS fragment ions. False discovery rate (FDR) of peptides was set to less than 1%. Three biological replicates were used for analysis.

### Measurement of metabolomics by UHPLC-MS/MS Analysis

Nine-day-old *Arabidopsis* wild type *Col_0* seedlings were treated by added 15 μL of 1 M GSNO into 100 mL 1/2 MS liquid medium, seedlings were grown at 22 °C. Samples were collected at 0, 6, and 24 h post-treatment. Five-day-old wheat *(Triticum aestivum* vs. Fielder) seedlings added 15 μL of 0.1 M GSNO into 10 mL liquid medium, seedling were grown at 22 °C, Samples were collected at 0, 6, and 24 h post-treatment.

### Untargeted metabolomics

Samples were weighed (150 mg) and mixed with 1 mL of 70% methanol solution. The mixture was vortexed for 30 s, left to stand at low temperature (4 °C) for 5 min, and then vortexed again. After overnight extraction at 4 °C, the mixture was centrifuged at 12,000 × g for 15 min at 4 °C. The supernatant was filtered through a 0.22-μm membrane filter, and the filtrate was further centrifuged at 12,000 g/min for 20 min at 4 °C. The final supernatant was collected for untargeted metabolomics analysis using LC-MS/MS.

Untargeted metabolomics was performed using a Vanquish F UHPLC system (Thermo Fisher Scientific, Germany) coupled with an Orbitrap Exploris 240 mass spectrometer (Thermo Fisher Scientific, Germany) equipped with a Heated Electrospray Ionization (HESI) by operating both in positive and negative ion modes. Samples were injected onto an ACQUITY UPLC HSS T3 column (2.1 mm × 150 mm, 1.8 μm) coupled with a VanGuard pre-column of the same stationary phase (2.1 mm × 5 mm, 1.8 μm). The mobile phase consisted of 0.1% formic acid in water (Phase A) and 0.1% formic acid in acetonitrile (Phase B). The flow rate was set at 0.3 mL/min with a column temperature of 40 °C, and the injection volume was 5 μL. The solvent gradient was set as follows: 0-2.5 min, 3-10% B; 2.5-6.0 min, 10-12% B; 6.0-10.0 min, 12-23% B; 10.0-12.4 min, 23-40% B; 12.4-20.0 min, 40-80% B; 20.0-22.0 min, 80-100% B; 22.0-25.0 min, maintained at 100% B; 25.0-25.1 min, 100-3% B; 25.1-30.0 min, maintained at 3% B. The major mass spectrometer parameters were set as below: data was acquired in full-scan combined with data-dependent secondary mass spectrometry (full scan + ddMS) mode; full scan mass range, *m/z* 80-,1200; spray voltage, 3,500 V for positive ion mode and 3,200 V for negative ion mode; sheath gas, 35 arb; auxiliary gas, 15 arb; ion transfer tube temperature, 350, vaporizer temperature, 350. Higher-energy collisional dissociation (HCD) was adopted for MS fragmentation with normalized collision energies of 20%, 40% and 60%.

### Untargeted metabolomics data analysis

The untargeted data were imported into Compound Discoverer software for peak extraction, alignment, compound detection and grouping, missing value imputation, SERRF-based QC correction, and background ion flagging. Compound identification was performed by molecular formula prediction and database searching against the online mzCloud and ChemSpider databases, as well as the local mzVault and Mass Lists databases. Finally, the identification results were filtered using the thresholds of mzCloud Best Match ≥ 60 and mzVault Best Match ≥ 70. Metabolite trends were classified based on the fold changes of 6 h vs. 0 h (r_1_) and 24 h vs. 6 h (r_2_). Metabolites were assigned to “Down” when r_1_ < 1, and r_2_ < 1, “Up” when r_1_ > 1.1 and r_2_ > 1.1, “Stable” when r_1_ < 1.1 and r_2_ < 1.1, and “Fluctuation” otherwise. The metabolites from the untargeted metabolomics analysis were mapped to biological pathways and subjected to pathway enrichment and classification analysis using the Pathway Analysis module in MetaboAnalyst.

### targeted metabolomics (GSH)

Samples were weighed (50 mg) and mixed with 1 mL of 0.1% formic acid in water solution. The mixture was vortexed for 30 s, sonicated at low temperature (4 °C) for 20 min, and then extracted at 4 °C. The mixture was centrifuged at 12,000 g/min for 15 min under 4 °C. The supernatant was filtered through a 0.22-μm membrane filter and diluted to 5 mg/mL, and the diluted extract was centrifuged again at 12,000 g/min for 20 min under 4 °C. The final supernatant was collected for targeted metabolomics analysis using LC-MS/MS. The detection were preformed on a Vanquish UHPLC system (Thermo Fisher Scientific, Germany) coupled with a TSQ Altis MS/MS system (Thermo Fisher Scientific, Germany) equipped with a Heated Electrospray Ionization (H-ESI) by operating both in positive and negative ion modes. Samples were injected onto an Acquity UPLC@HSS T3 Column (1.8 µm, 2.1 × 150 mm). The mobile phase consisted of 0.1% formic acid-water (Phase A) and 0.1% formic acid in acetonitrile (Phase B). The flow rate was set at 0.2 mL/min with a column temperature of 25 °C, and the injection volume was 2 μL. The solvent gradient was set as follows: 0-1 min, 0% B; 1-2 min, 0-3% B; 2-5 min, 3-90% B; 5-11 min, 90% B; 11-12.5 min, 90-60% B; 12.5-13 min, 60-0% B. The major mass spectrometer parameters were set as below: 4000 V for positive ion mode; sheath gas: 35 arb; auxiliary gas: 10 arb; ion transfer tube temperature: 350 °C, vaporizer temperature: 350 °C. The MRM transitions for reduced glutathione (GSH) were m/z 307.883, and 178.800, with collision energies of 12.23 V, and RF lens voltage of 48 V. The linear equation was y =-140879 + 20504.5 x, R^2^ = 0.9994.

### Laser scanning confocal imaging

Using the Beyotime S0020S Kit, take wheat or *Arabidopsis* root tips (about 1 cm), soak them in 10 μM of the cell-permeable fluorescent probe DAF-FM DA, and leave them in the dark for 30 min. Then rinse the roots three times with buffer. Observe using an upright fluorescence microscope, with excitation at 488 nm and emission at 495-575 nm (ZEISS Axio Imager M2), and record the root fluorescence under brightfield and darkfield conditions.

### Gene Ontology analysis

GO analysis and KEGG were performed using DAVID (2025) Bioinformatics Resources. Pricicpal component analysis (PCA) of all sample using the prcomp function from the stats package in R (v 4.5.2). Consensus sequences of S-nitrosylated peptides were predicted using the rmotifx method^17^. Residue heights were scaled relative to their statistical significance, calculated as the log-odds of the binomial probability of overrepresentation versus underrepresentation against a defined background model. To enhance visual interpretability, residues within each position were reordered such that the most statistically significant residues are positioned closest to the x-axis. The statistical significance threshold was determined using a Bonferroni correction for multiple testing, calculated as 0.05/ 15-1 20, corresponding to the specific window size and residue alphabet used in this analysis.

### Protein expression and purification

For recombinant protein expression, the coding sequence of AtGR1 was subcloned into pGEX-6P1 vector to fuse a GST tag to the N-terminus of the target protein. The plasmids were transformed into *Escherichia coli* strain BL21 (DE3) RIL (Tsingke). Expression was induced at 16 °C overnight with 0.3 mM of IPTG. The recombinant proteins were purified with a prepackaged GST Sep Glutathione Agarose Resin column (cat# 202507ES10, Yeasen Biotechnology, Shanghai, China). The GST tag was removed by HRV 3C protease (cat# Z03092-250, GeneScript Biotechnology, Shanghai, China) before use in further activity assay.

For recombinant protein expression, the coding sequence of TaGR1-B was subcloned into pET-28a vector to fuse a His tag to the C-terminus of the target protein. The plasmids were transformed into *Escherichia coli* strain BL21 (DE3) RIL (Tsingke). Expression was induced at 16 °C overnight with 0.2 mM of IPTG. The recombinant proteins were purified with a prepackaged Ni Sepharose 6 Fast Flow column (cat# 17531806, GE Healthcare). The primers used for vector construction are listed as below: pET-28a-TaGR1-B F: GGTGGACAGCAAATGGGTCGCATGCTCAAGGACGGGGAGG pET-28a-TaGR1-B R: CGGAGCTCGAATTCGGATCCCAAGTTCGTCTTTGGCTTG pGEX-6P1-AtGR1 F: AGTTCTGTTCCAGGGGCCCCTGATGGCGAGGAAGATGCTTG pGEX-6P1-AtGR1 R: GGGAATTCCGGGGATCCTCATAGATTTGTCTTAGGTTTGG

### Immunoblot Assay

For in vitro S-nitrosylation assay by immunoblotting, approximately 100 μg recombinant GR protein was incubated with 1 mM GSNO or GSH at room temperature in the dark for 60 min, and excess GSNO or GSH was removed using a 10 kDa filter. The sample was blocked as described above and dissolved in 100 μL of HENS buffer, followed by adding 5 μL of 200 mM sodium ascorbate and 25 μL of 4 mM biotin-maleimide. After labeling for 90 min at room temperature with shaking, the proteins were washed two times with neutralization buffer (25 mM HEPES, pH 7.7, 100 mM NaCl, 1 mM EDTA, 0.5% Triton X-100) in 10 kDa filter to remove biotin-maleimide. The proteins were resolubilized in 1 mL neutralization buffer, and incubated with 30 μL of streptavidin beads at 4 °C overnight. After the incubation, the beads were washed five times with wash buffer (25 mM HEPES, pH 7.7, 600 mM NaCl, 1 mM EDTA, 0.5% Triton X-100). Eluted proteins were recovered with 30 μL elution buffer (250 mM HEPES, pH 7.7, 1 mM EDTA, 1% 2-mercaptoethanol) and subsequently subjected to SDS-PAGE analysis. Primary antibodies used in this study are as follows: anti-GR1 (Agrisera, AS06181), anti-His.

### GR1 activity analysis

For GR1 activity, approximately 15 μg of recombinant GR protein was incubated with 1 mM GSNO or GSH at room temperature in the dark for 60 min, and excess GSNO or GSH was removed using 10 kDa filter. The enzyme activity was estimated using a Glutathione Reductase Assay Kit with DTNB [5,5-dithio-bis-(2-nitrobenzoic acid)] colorimetry (cat#S0055; Beyotime Biotechnology, Shanghai, China). The absorbance at 412 nm was continuously recorded every 2 min for 10 min at 25 °C.

## DATA AVAILABILITY

The mass spectrometry proteomics data have been deposited to the Proteome Xchange Consortium via the PRIDE partner repository with the dataset identifier PXD081361. Source data are provided with this paper.

## ACKNOWLEDGEMENTS

We thank Proteomics and Mass Spectrometry Centre in Peking University Institute of Advanced Agricultural Sciences for the LC-MS/MS services.

This work was supported by National Key R & D Program of China (Grant No. 2023YFD1202100) and the Shandong Provincial Natural Science Foundation (SYS202206).

## AUTHOR CONTRIBUTIONS

These authors contributed equally to the study: Ruixue Ji, Ning Yang.

## CORRESPONDING AUTHORS

Correspondence to Guochen Qin.

## SUPPLEMENTARY INFORMATION

Supplementary Information: should be combined and supplied as a separate file, preferably in PDF format.

## Supplementary Figures

**Supplementary Figure S1**. Workflow of F-MNPs. (Supplementary Fig. S1)

**Supplementary Figure S2.** PCA analysis, and confocal image from GSNO treatment for *Arabidopsis* seedling. (Supplementary Fig. S2)

**Supplementary Figure S3**. Motif analysis from GSNO treatment for *Arabidopsis* seedling. (Supplementary Fig. S3)

**Supplementary Figure S4**. GO analysis from GSNO treatment for *Arabidopsis* seedling. (Supplementary Fig. S4)

Supplementary Figure S5. Untargeted metabolomic of *Arabidopsis* seedling under GSNO treatment. (Supplementary Fig. S5)

**Supplementary Figure S6**. Analysis of GSNO treatment wheat Fielder within 1 h. (Supplementary Fig. S6)

**Supplementary Figure S7**. Metabolomic changes in wheat after GSNO treatment. (Supplementary Fig. S7)

**Supplementary Figure S8**. GSH abundance in *Arabidopsis thaliana* and wheat upon GSNO treatment. (Supplementary Fig. S8)

**Supplementary Figure S9**. Schematic diagram of gene editing and phenotypic analysis. (Supplementary Fig. S9)

## Supplementary Tables

**Supplementary Table S1**. TEM-EXD Element Analysis. (Supplementary Table. S1)

**Supplementary Table S2**. S-nitrosylated peptides enrichment by different amount of Fe_3_O_4_@SiO_2_@TFS. (Supplementary Table. S2)

**Supplementary Table S3**. Comparison of the results obtained by F-MNPs methods in this study with data from the nGF literature. (Supplementary Table. S3)

**Supplementary Table S4.** Identified modification in 73 carbon metabolism-associated proteins from *Arabidopsis* under GSNO treatment 0.5 h. (Supplementary Table. S4)

**Supplementary Table S5**. S-nitrosylated proteins involved in GSH biosynthesis and metabolism in *Arabidopsis* and wheat following GSNO treatment. (Supplementary Table. S5)

**Supplementary Table S6**. Intensity of S-nitrosylated peptides containing Cys 73 of *Arabidopsis* GR1 and Cys 71 of wheat GR1following GSNO treatment. (Supplementary Table. S6)

## Supplementary Datas

**Supplementary Data 1.** S-nitrosylated peptides enriched from *Col_0* under GSNO treatment.

**Supplementary Data 2.** Untargeted metabolomic of *Col_0* under GSNO treatment.

**Supplementary Data 3.** S-nitrosylated peptides enriched from Fielder under GSNO treatment.

**Supplementary Data 4.** Untargeted metabolomic of Fielder under GSNO treatment.

## 1. Supplementary Figures

**Supplementary Fig. S1.**
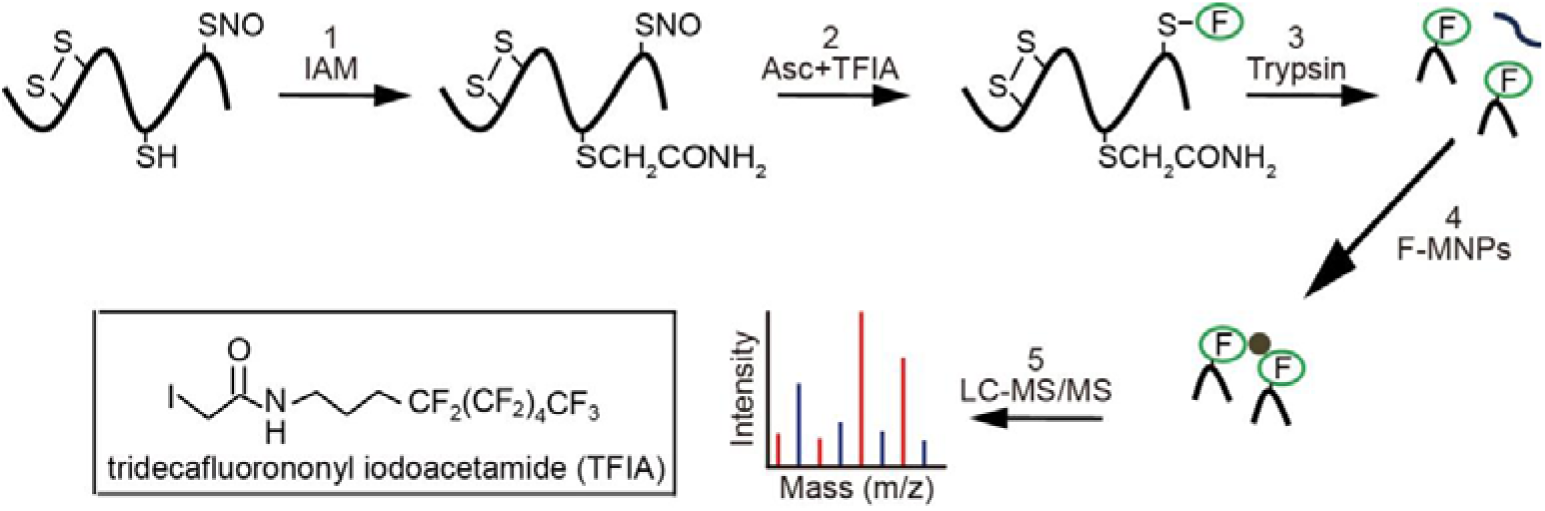
Reaction schema for FAT-switch approach to label S-nitrosylated proteins and enrich target peptides. Step 1, free cysteine thiols were blocked by IAA. In step 2, the S-nitrosylated cysteines were specifically reduced by a low concentration of ascorbate and substituted by a sulfhydryl reactive fluorous tag reagent. Step 3, proteins were digested by trypsin. After that, Fe_3_O_4_@SiO_2_@TFS nanoparticle were used to capture fluorous-tag labeling peptides and remove high hydrophilic peptide by a low ACN concentration of washing buffer (Step 4). Finally, fluorous labeled peptides were identified by LC-MS/MS (Step 5).

**Supplementary Fig. S2.**
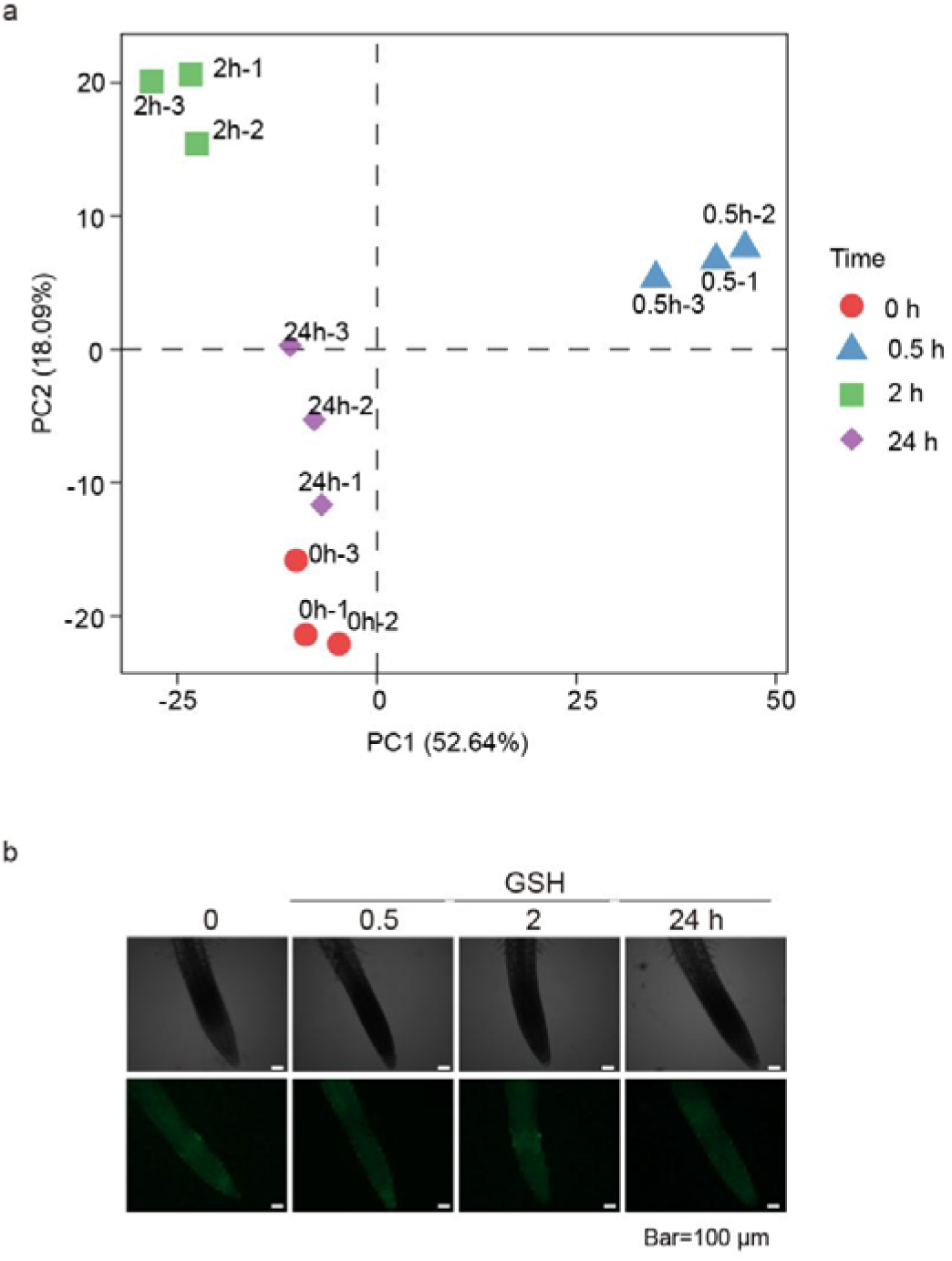
a. The principal component analysis (PCA) of the global S-nitrosylated peptides from *Arabidopsis* seedling under GSNO treatment. **b** Confocal microscopy of 9-day-old *Arabidopsis* root of GSH treatment using DAF-FM to NO distribution. Image brightness and contrast were uniformly adjusted across all panels using ImageJ for display purposes.

**Supplementary Fig. S3.**
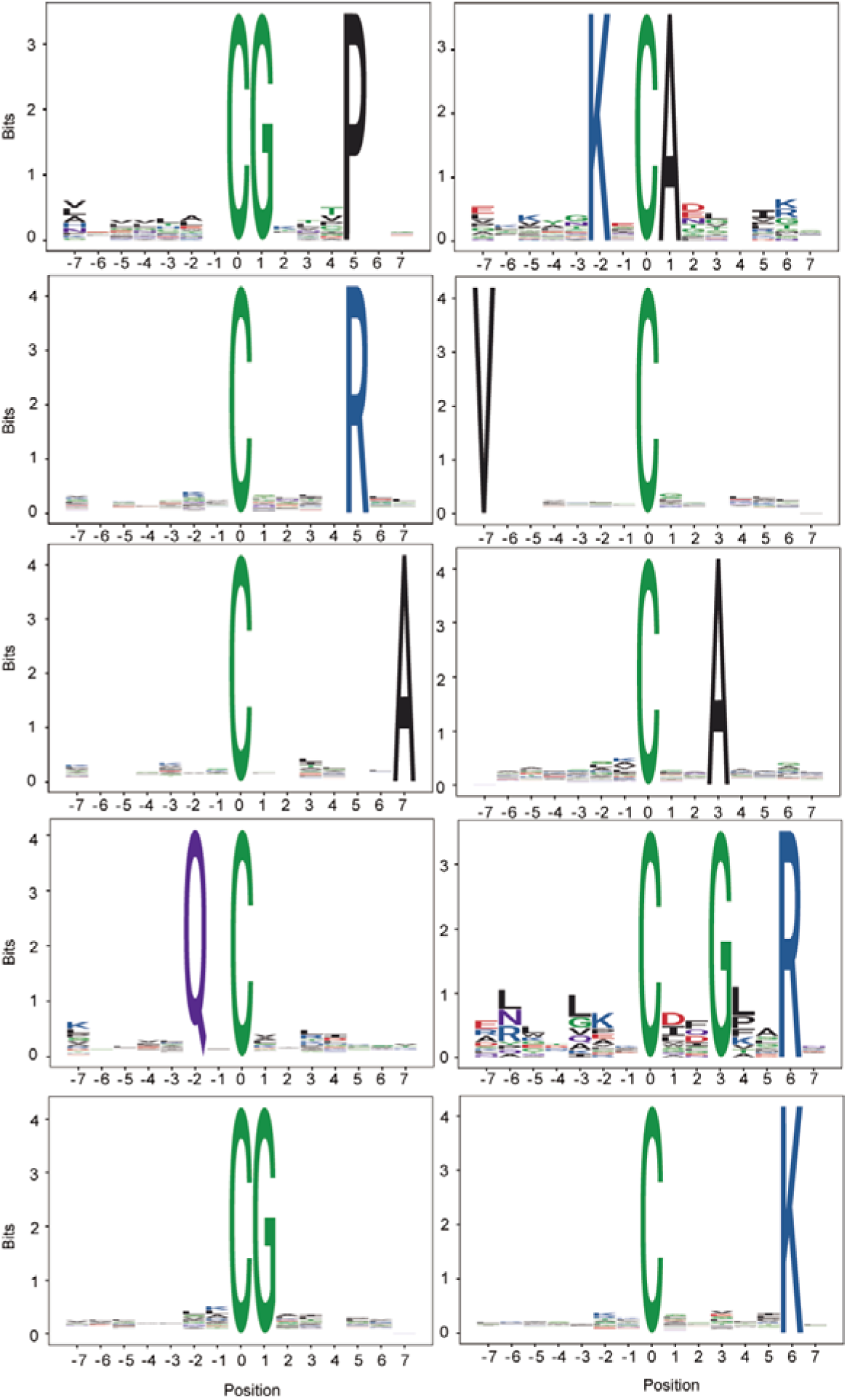
Motif analysis of 1171 S-nitrosylation sites which were up-regulated at 0.5 h post-GSNO treatment for *Arabidopsis* seedling. Over-representation motif of Cys-SNO sequence windows (-7 to +7 residues) constructed by rmotifx.

**Supplementary Fig. S4.**
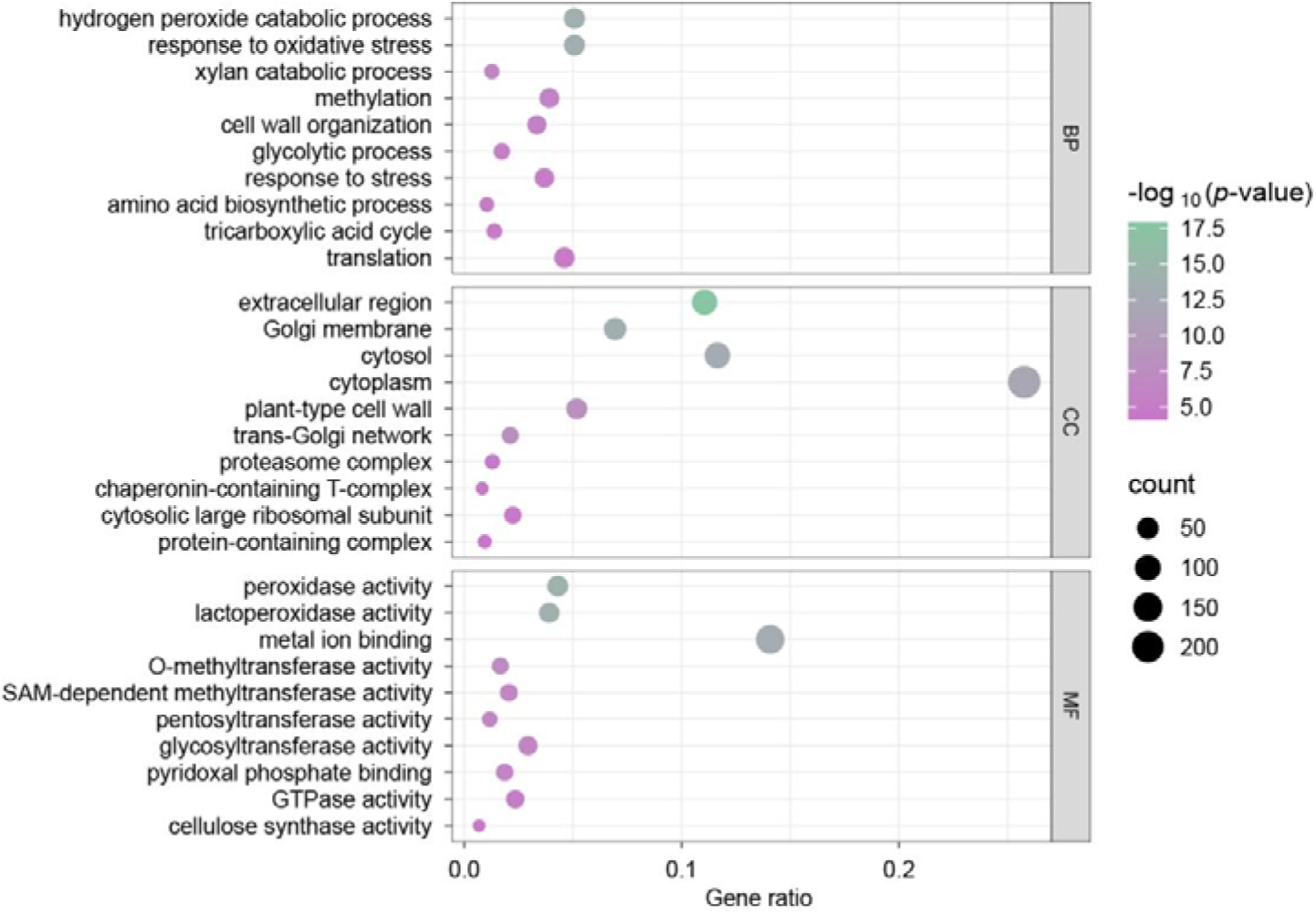
The top 10 significantly enriched Gene Ontology (GO) terms for biological process (BP), cellular component (CC), and molecular function (MF) categories derived from the *Arabidopsis* proteins differentially expressed under GSNO treatment for 0.5 h.

**Supplementary Fig. S5.**
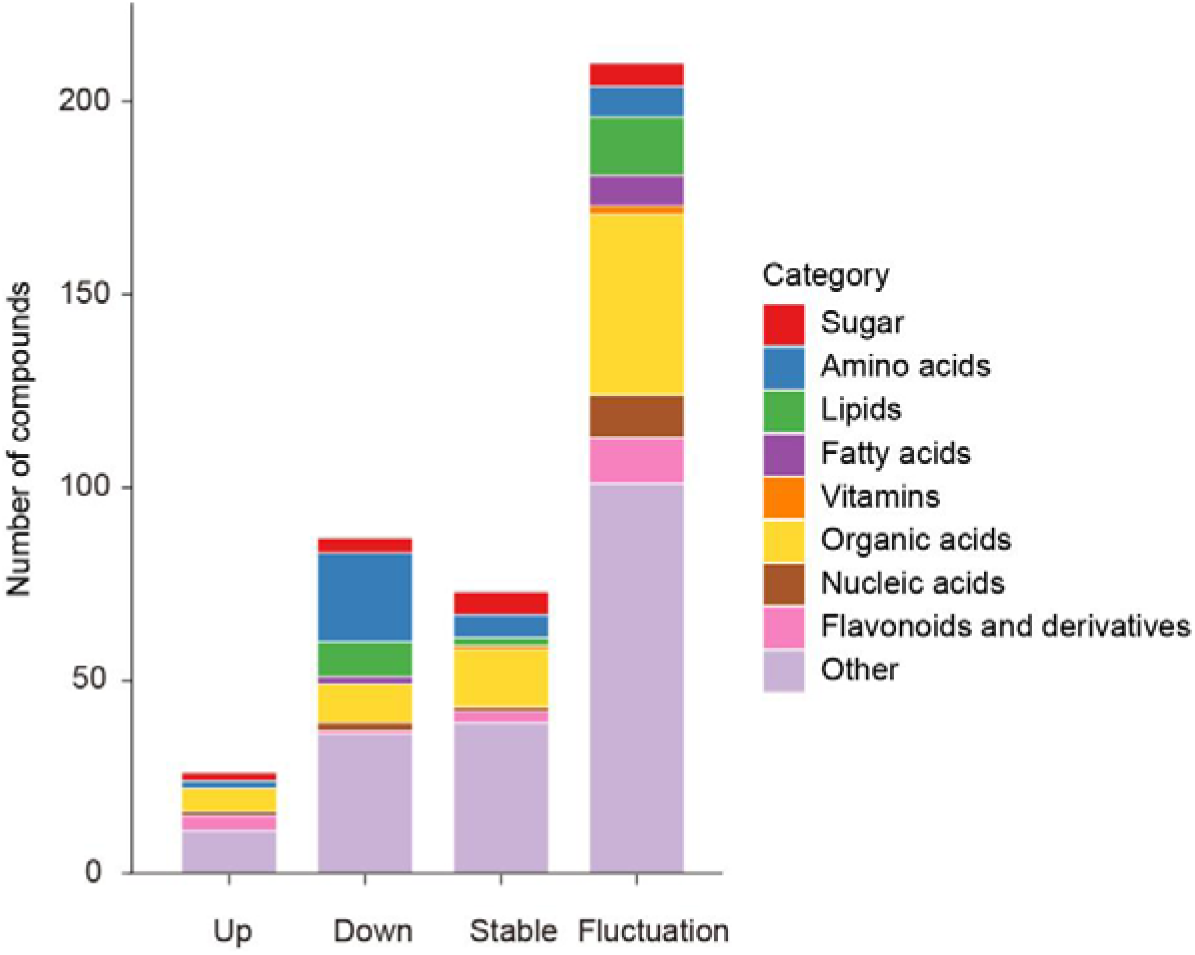
**Untargeted metabolomic of *Arabidopsis* seedling under GSNO treatment**. Stacked bar chart showing the classification and change trends of compounds assigned to the untargeted compounds across different dynamic change patterns.

**Supplementary Fig. S6.**
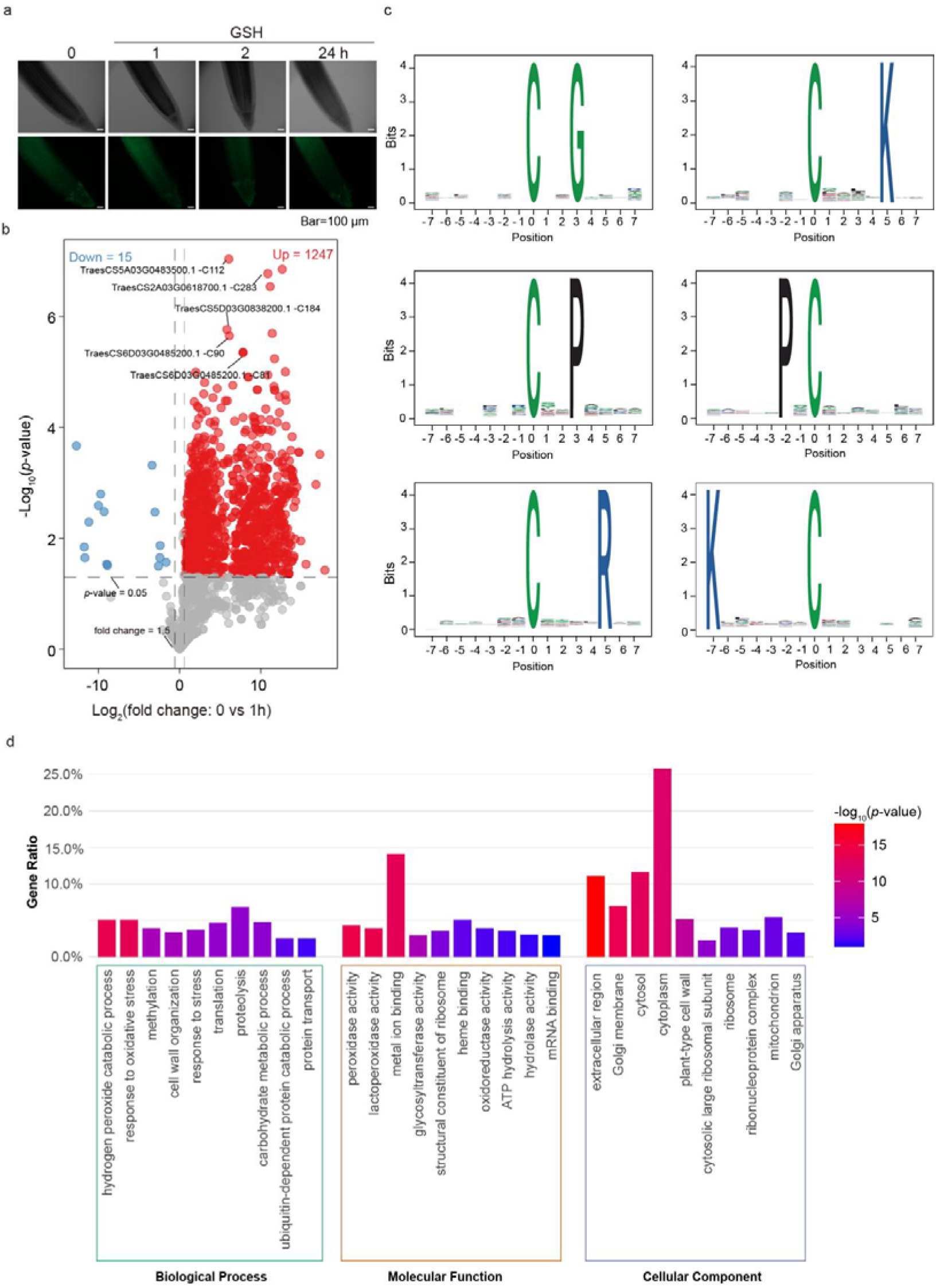
a. Volcano plot of S-nitrosylation site intensity in 1 h post-GSNO treatment. Red dots (n = 1247) represent S-nitrosylated sites that were up-regulated at 1 h post-GSNO treatment, blue dots (n = 15) represent S-nitrosylated sites that were down-regulated at 1 h post-GSNO treatment (|fold change| > 1.5, *p* < 0.05, two-tailed unpaired *t* test). **b** Motif analysis of 942 S-nitrosylation sites which up-regulated from fielder at 1 h-post GSNO treatment. Over representation motif of Cys-SNO sequence windows (-7 to +7 residues) constructed by rmotifx. **c** Confocal microscopy of 5-day-old Fielder root of GSH treatment using DAF-FM to NO distribution. Image brightness and contrast were uniformly adjusted across all panels using ImageJ for display purposes. **d** The top 10 significantly enriched GO for biological process (BP), cellular component (CC), and molecular function (MF) categories derived which from GSNO treatment wheat Fielder within 1 h.

**Supplementary Fig. S7.**
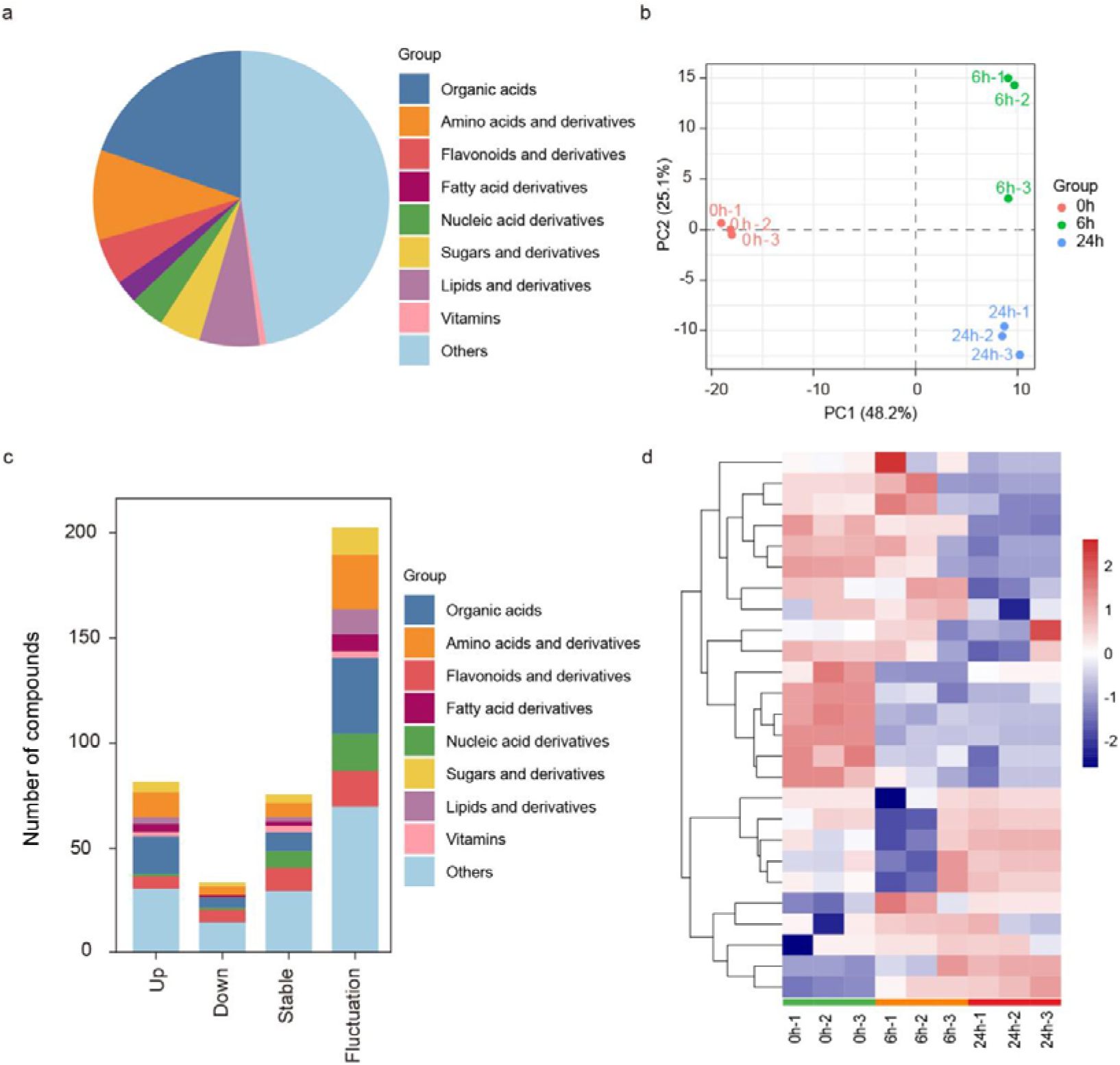
Metabolomic changes in wheat after GSNO treatment. **a** Statistics of untargeted compounds across different classification from the Fielder under GSNO treatment for 0 h, 6 h, and 24 h. **b** Pricipal component analysis of metabolites from the Fielder under GSNO treatment for 0 h, 6 h, and 24 h. **c** Stacked bar chart showing the classification and change trends of compounds assigned to the untargeted compounds across different dynamic change patterns. **d** The heatmap of compounds on carbon metabolism pathway from the Fielder under GSNO treatment for 0 h, 6 h, and 24 h.

**Supplementary Fig. S8.**
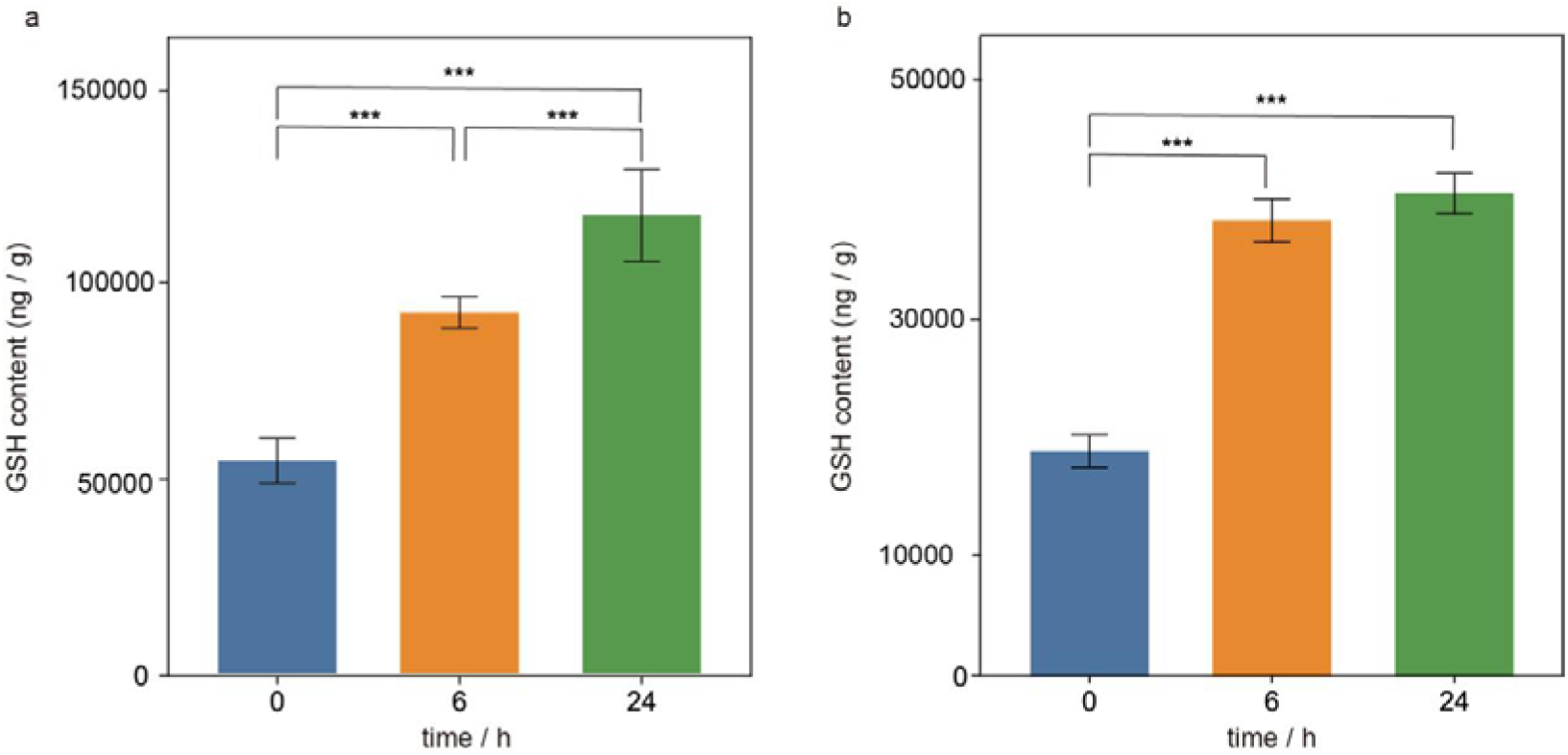
a. GSH abundance in *Arabidopsis thaliana* upon GSNO treatment. **B** GSH abundance in wheat upon GSNO treatment. Error bars, SD (n = 3 biological replicates), *** *p* < 0.005, one-way ANOVA.

**Supplementary Fig. S9.**
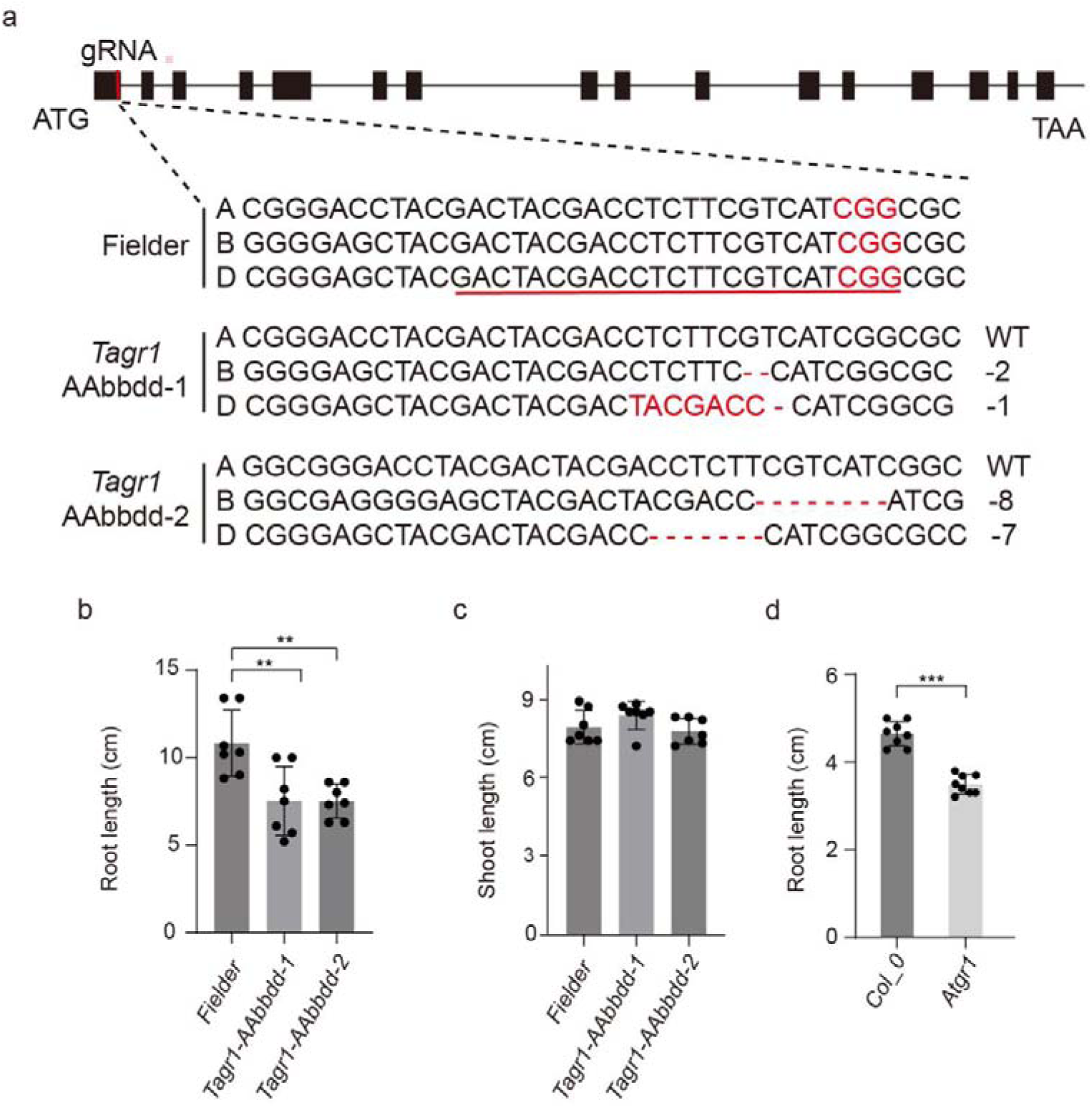
a. Different types of CRISPR/Cas9-induced mutations in TaGR1 homologs. “-”indicate the deletion and the numbers indicate the length of the deletion. **b** Root length of Fielder (wild type), *Tagr1-AAbbdd-1* and *Tagr1-AAbbdd-2* plants. Error bars, SD (n = 7 seedlings). ** *p* < 0.01, one-way ANOVA. **c** Shoot length of Fielder (wild type), *Tagr1-AAbbdd-1* and *Tagr1-AAbbdd-2* plants. Error bars, SD (n = 7 seedlings). **d** Root length of *Col_0* (wild type) and *Atgr1* plants. Error bars, SD (n = 8 seedlings). *** *p* < 0.005, one-way ANOVA.

## 2. Supplementary Tables

**Supplementary Table S1.**
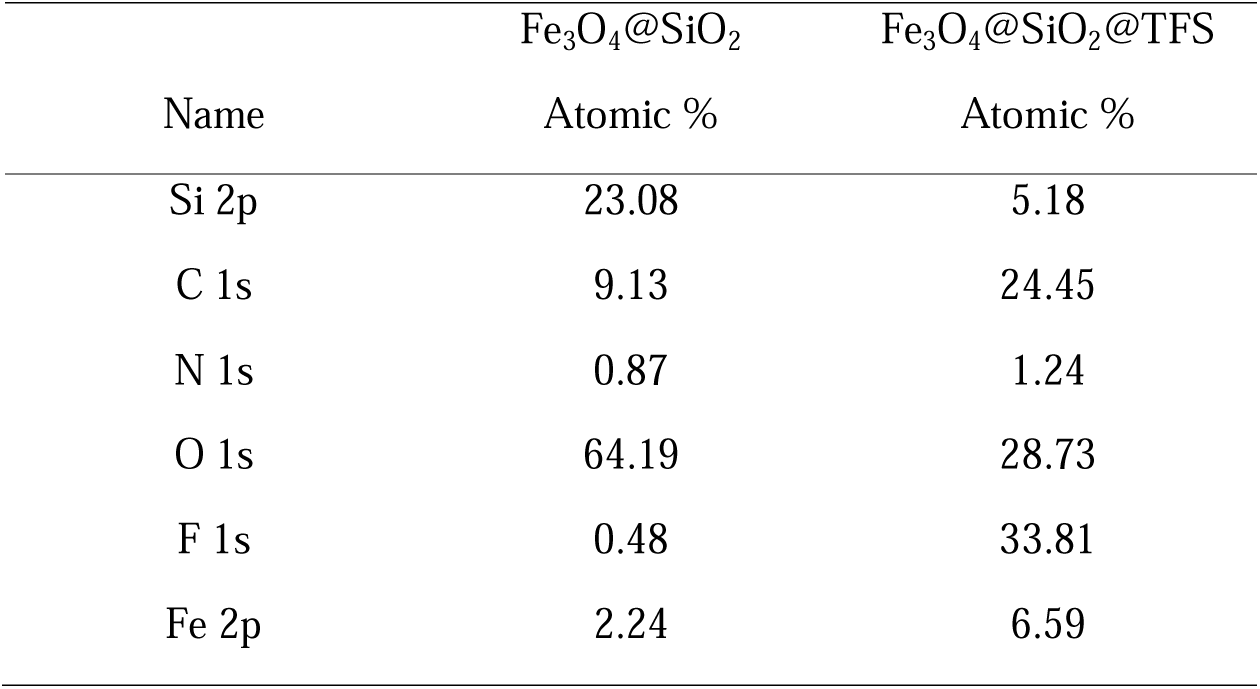
TEM-EXD Element Analysis.

|  | Fe<sub>3</sub>O<sub>4</sub>@SiO<sub>2</sub> | Fe<sub>3</sub>O<sub>4</sub>@SiO<sub>2</sub>@TFS |
|---|---|---|
| Name | Atomic % | Atomic % |
| Si 2p | 23.08 | 5.18 |
| C 1s | 9.13 | 24.45 |
| N 1s | 0.87 | 1.24 |
| O 1s | 64.19 | 28.73 |
| F 1s | 0.48 | 33.81 |
| Fe 2p | 2.24 | 6.59 |

**Supplementary Table S2.** S-nitrosylated peptides enrichment by different amount of Fe_3_O_4_@SiO_2_@TFS.

|  | 5 mg | 10 mg | 15 mg |
|---|---|---|---|
| Rep 1 | 1530 | 1767 | 1040 |
| Rep 2 | 1480 | 1854 | 1374 |
| Rep 3 | 1395 | 1808 | 1402 |
| Average ± SD | 1468.33 ± 68.25 | 1810.67 ± 43.52 | 1272.00 ± 201.41 |
| enrichment efficiency | 12.11% | 14.05% | 9.43% |

**Supplementary Table S3.** Comparison of the results obtained by F-MNPs methods in this study with data from the nGF literature^12^.

|  | F-MNPs | nGF |
|---|---|---|
| Rep 1 | 1767 | 1920 |
| Rep 2 | 1854 | 1760 |
| Rep 3 | 1808 | 1641 |
| Average ± SD | 1809.67 ± 43.52 | 1773.67 ± 140.00 |
| CV | 2.41% | 7.89% |

**Supplementary Table S4.**
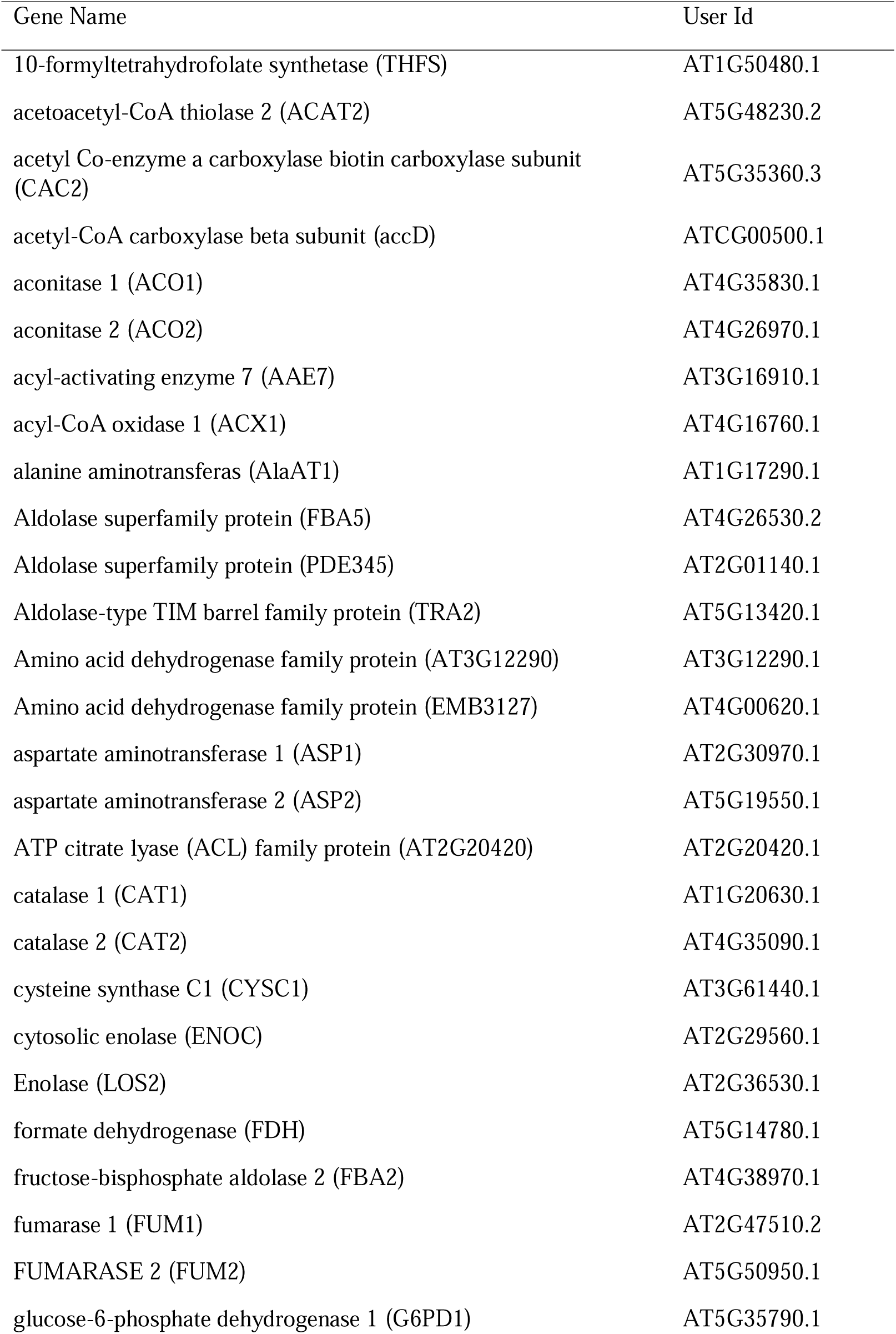
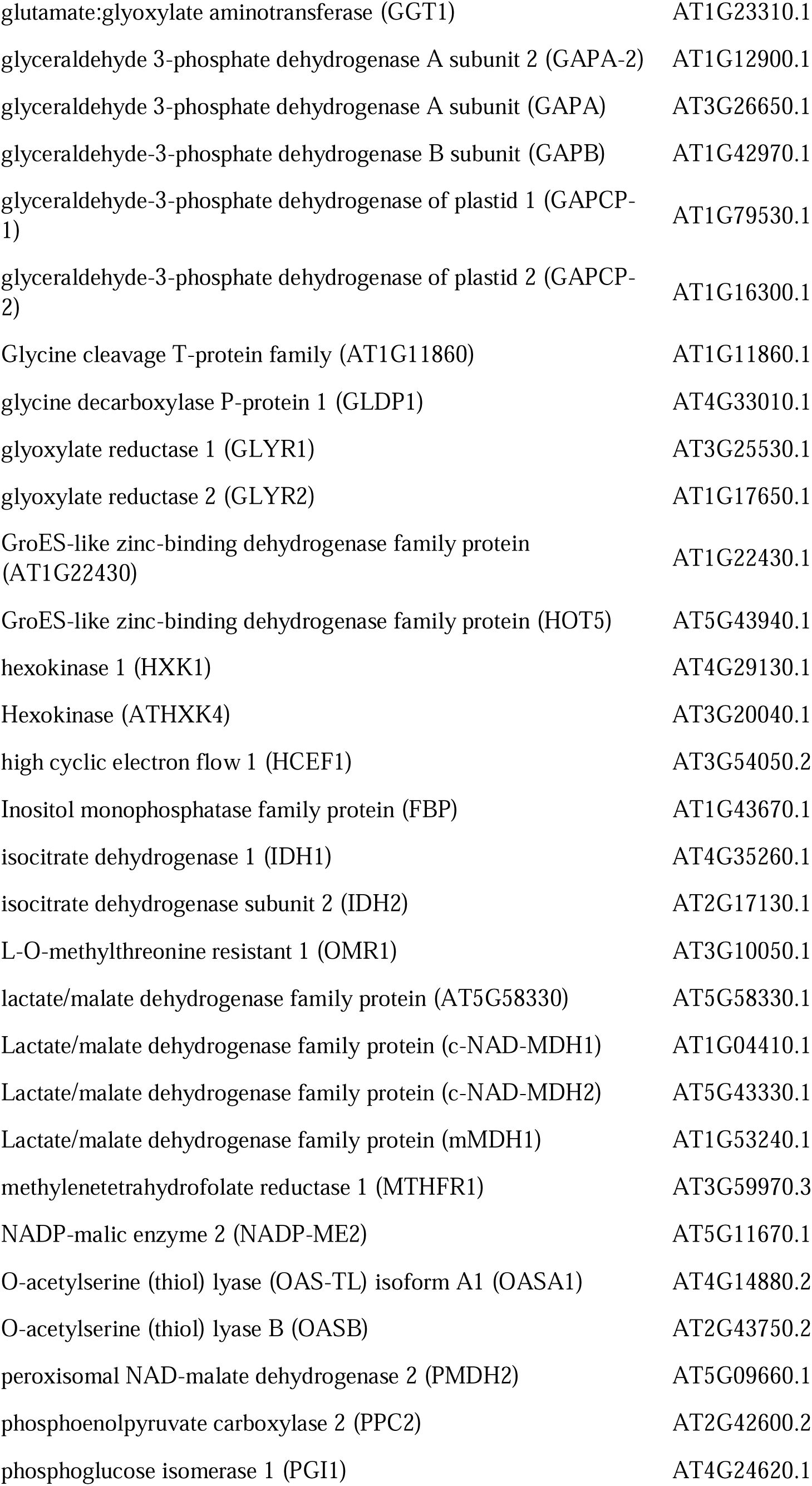
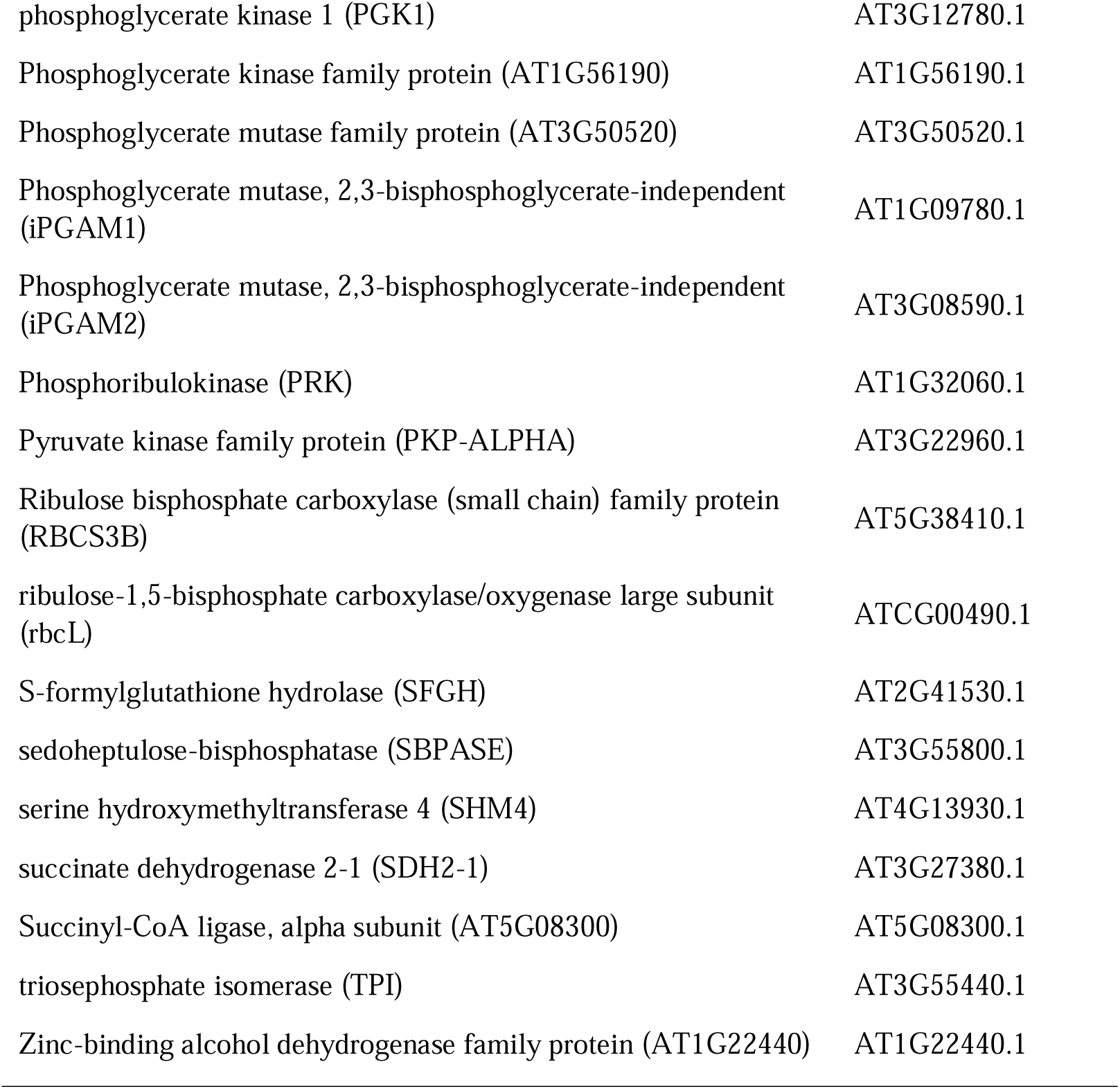
Identified modifications in carbon metabolism-associated proteins from *Arabidopsis* under GSNO treatment 0.5 h.

| Gene Name | User Id |
|---|---|
| 10-formyltetrahydrofolate synthetase (THFS) | AT1G50480.1 |
| acetoacetyl-CoA thiolase 2 (ACAT2) | AT5G48230.2 |
| acetyl Co-enzyme a carboxylase biotin carboxylase subunit (CAC2) | AT5G35360.3 |
| acetyl-CoA carboxylase beta subunit (accD) | ATCG00500.1 |
| aconitase 1 (ACO1) | AT4G35830.1 |
| aconitase 2 (ACO2) | AT4G26970.1 |
| acyl-activating enzyme 7 (AAE7) | AT3G16910.1 |
| acyl-CoA oxidase 1 (ACX1) | AT4G16760.1 |
| alanine aminotransferas (AlaAT1) | AT1G17290.1 |
| Aldolase superfamily protein (FBA5) | AT4G26530.2 |
| Aldolase superfamily protein (PDE345) | AT2G01140.1 |
| Aldolase-type TIM barrel family protein (TRA2) | AT5G13420.1 |
| Amino acid dehydrogenase family protein (AT3G12290) | AT3G12290.1 |
| Amino acid dehydrogenase family protein (EMB3127) | AT4G00620.1 |
| aspartate aminotransferase 1 (ASP1) | AT2G30970.1 |
| aspartate aminotransferase 2 (ASP2) | AT5G19550.1 |
| ATP citrate lyase (ACL) family protein (AT2G20420) | AT2G20420.1 |
| catalase 1 (CAT1) | AT1G20630.1 |
| catalase 2 (CAT2) | AT4G35090.1 |
| cysteine synthase C1 (CYSC1) | AT3G61440.1 |
| cytosolic enolase (ENOC) | AT2G29560.1 |
| Enolase (LOS2) | AT2G36530.1 |
| formate dehydrogenase (FDH) | AT5G14780.1 |
| fructose-bisphosphate aldolase 2 (FBA2) | AT4G38970.1 |
| fumarase 1 (FUM1) | AT2G47510.2 |
| FUMARASE 2 (FUM2) | AT5G50950.1 |
| glucose-6-phosphate dehydrogenase 1 (G6PD1) | AT5G35790.1 |
| glutamate:glyoxylate aminotransferase (GGT1) | AT1G23310.1 |
| glyceraldehyde 3-phosphate dehydrogenase A subunit 2 (GAPA-2) | AT1G12900.1 |
| glyceraldehyde 3-phosphate dehydrogenase A subunit (GAPA) | AT3G26650.1 |
| glyceraldehyde-3-phosphate dehydrogenase B subunit (GAPB) | AT1G42970.1 |
| glyceraldehyde-3-phosphate dehydrogenase of plastid 1 (GAPCP-1) | AT1G79530.1 |
| glyceraldehyde-3-phosphate dehydrogenase of plastid 2 (GAPCP-2) | AT1G16300.1 |
| Glycine cleavage T-protein family (AT1G11860) | AT1G11860.1 |
| glycine decarboxylase P-protein 1 (GLDP1) | AT4G33010.1 |
| glyoxylate reductase 1 (GLYR1) | AT3G25530.1 |
| glyoxylate reductase 2 (GLYR2) | AT1G17650.1 |
| GroES-like zinc-binding dehydrogenase family protein (AT1G22430) | AT1G22430.1 |
| GroES-like zinc-binding dehydrogenase family protein (HOT5) | AT5G43940.1 |
| hexokinase 1 (HXK1) | AT4G29130.1 |
| Hexokinase (ATHXK4) | AT3G20040.1 |
| high cyclic electron flow 1 (HCEF1) | AT3G54050.2 |
| Inositol monophosphatase family protein (FBP) | AT1G43670.1 |
| isocitrate dehydrogenase 1 (IDH1) | AT4G35260.1 |
| isocitrate dehydrogenase subunit 2 (IDH2) | AT2G17130.1 |
| L-O-methylthreonine resistant 1 (OMR1) | AT3G10050.1 |
| lactate/malate dehydrogenase family protein (AT5G58330) | AT5G58330.1 |
| Lactate/malate dehydrogenase family protein (c-NAD-MDH1) | AT1G04410.1 |
| Lactate/malate dehydrogenase family protein (c-NAD-MDH2) | AT5G43330.1 |
| Lactate/malate dehydrogenase family protein (mMDH1) | AT1G53240.1 |
| methylenetetrahydrofolate reductase 1 (MTHFR1) | AT3G59970.3 |
| NADP-malic enzyme 2 (NADP-ME2) | AT5G11670.1 |
| O-acetylserine (thiol) lyase (OAS-TL) isoform A1 (OASA1) | AT4G14880.2 |
| O-acetylserine (thiol) lyase B (OASB) | AT2G43750.2 |
| peroxisomal NAD-malate dehydrogenase 2 (PMDH2) | AT5G09660.1 |
| phosphoenolpyruvate carboxylase 2 (PPC2) | AT2G42600.2 |
| phosphoglucose isomerase 1 (PGI1) | AT4G24620.1 |
| phosphoglycerate kinase 1 (PGK1) | AT3G12780.1 |
| Phosphoglycerate kinase family protein (AT1G56190) | AT1G56190.1 |
| Phosphoglycerate mutase family protein (AT3G50520) | AT3G50520.1 |
| Phosphoglycerate mutase, 2,3-bisphosphoglycerate-independent (iPGAM1) | AT1G09780.1 |
| Phosphoglycerate mutase, 2,3-bisphosphoglycerate-independent (iPGAM2) | AT3G08590.1 |
| Phosphoribulokinase (PRK) | AT1G32060.1 |
| Pyruvate kinase family protein (PKP-ALPHA) | AT3G22960.1 |
| Ribulose bisphosphate carboxylase (small chain) family protein (RBCS3B) | AT5G38410.1 |
| ribulose-1,5-bisphosphate carboxylase/oxygenase large subunit (rbcL) | ATCG00490.1 |
| S-formylglutathione hydrolase (SFGH) | AT2G41530.1 |
| sedoheptulose-bisphosphatase (SBPASE) | AT3G55800.1 |
| serine hydroxymethyltransferase 4 (SHM4) | AT4G13930.1 |
| succinate dehydrogenase 2-1 (SDH2-1) | AT3G27380.1 |
| Succinyl-CoA ligase, alpha subunit (AT5G08300) | AT5G08300.1 |
| triosephosphate isomerase (TPI) | AT3G55440.1 |
| Zinc-binding alcohol dehydrogenase family protein (AT1G22440) | AT1G22440.1 |

**Supplementary Table S5.**
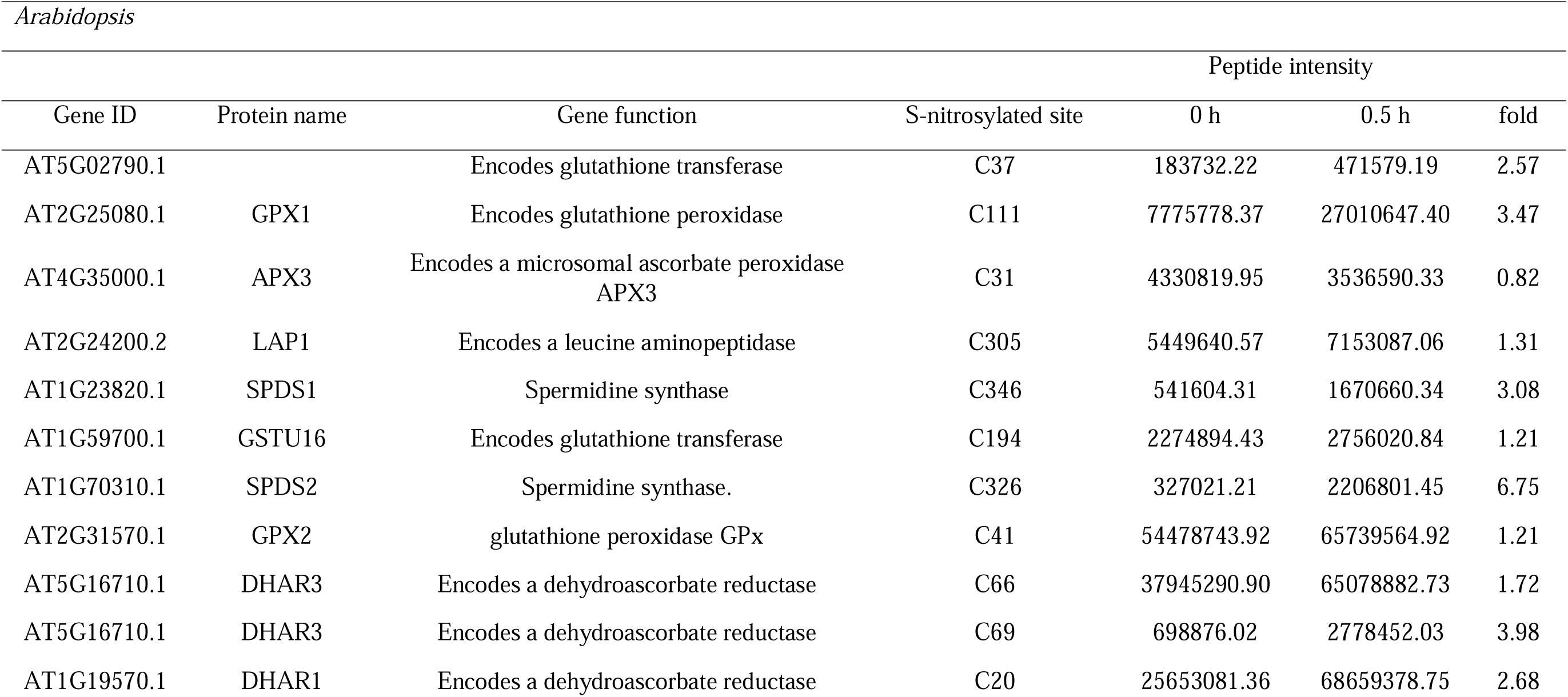
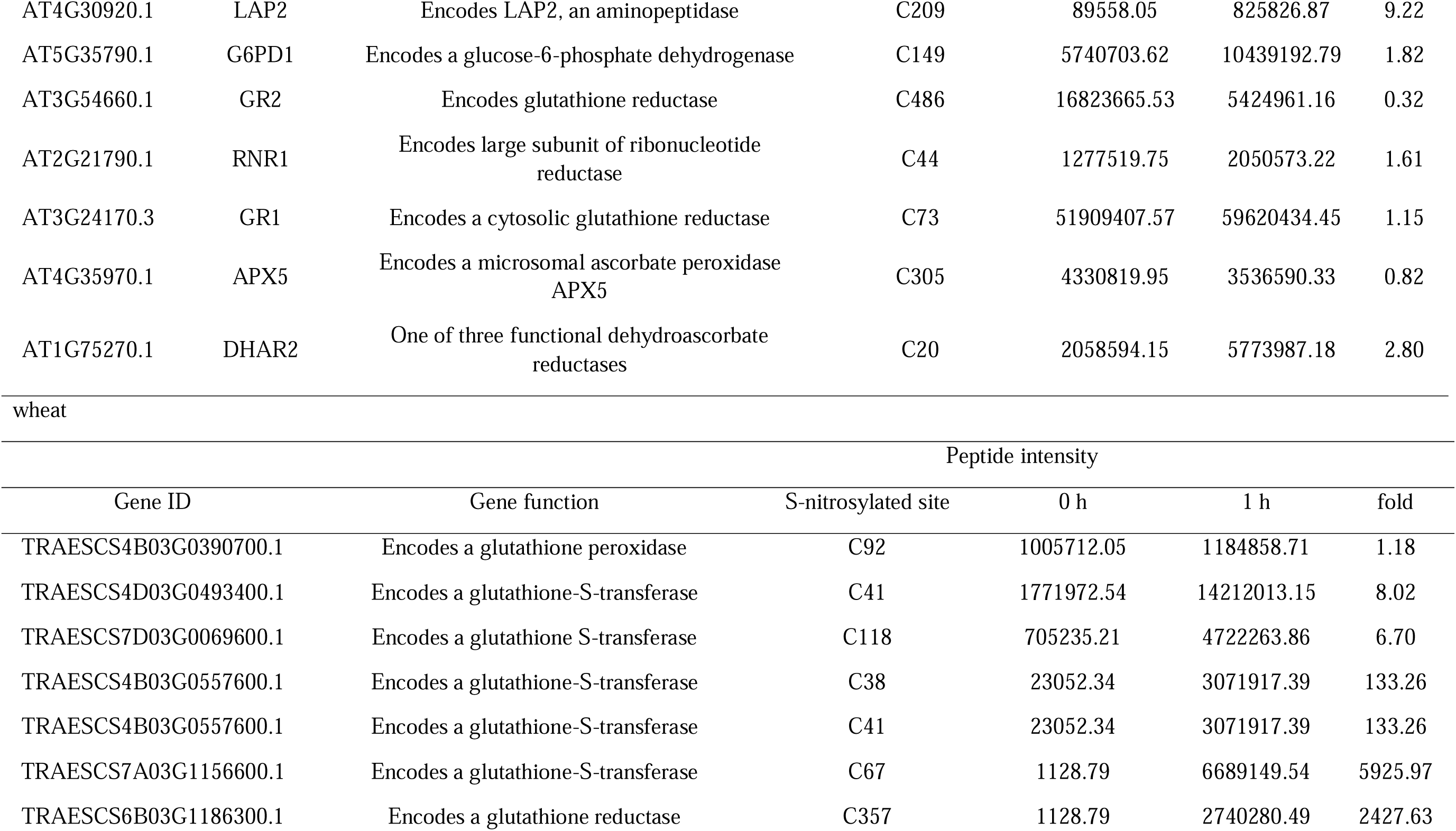
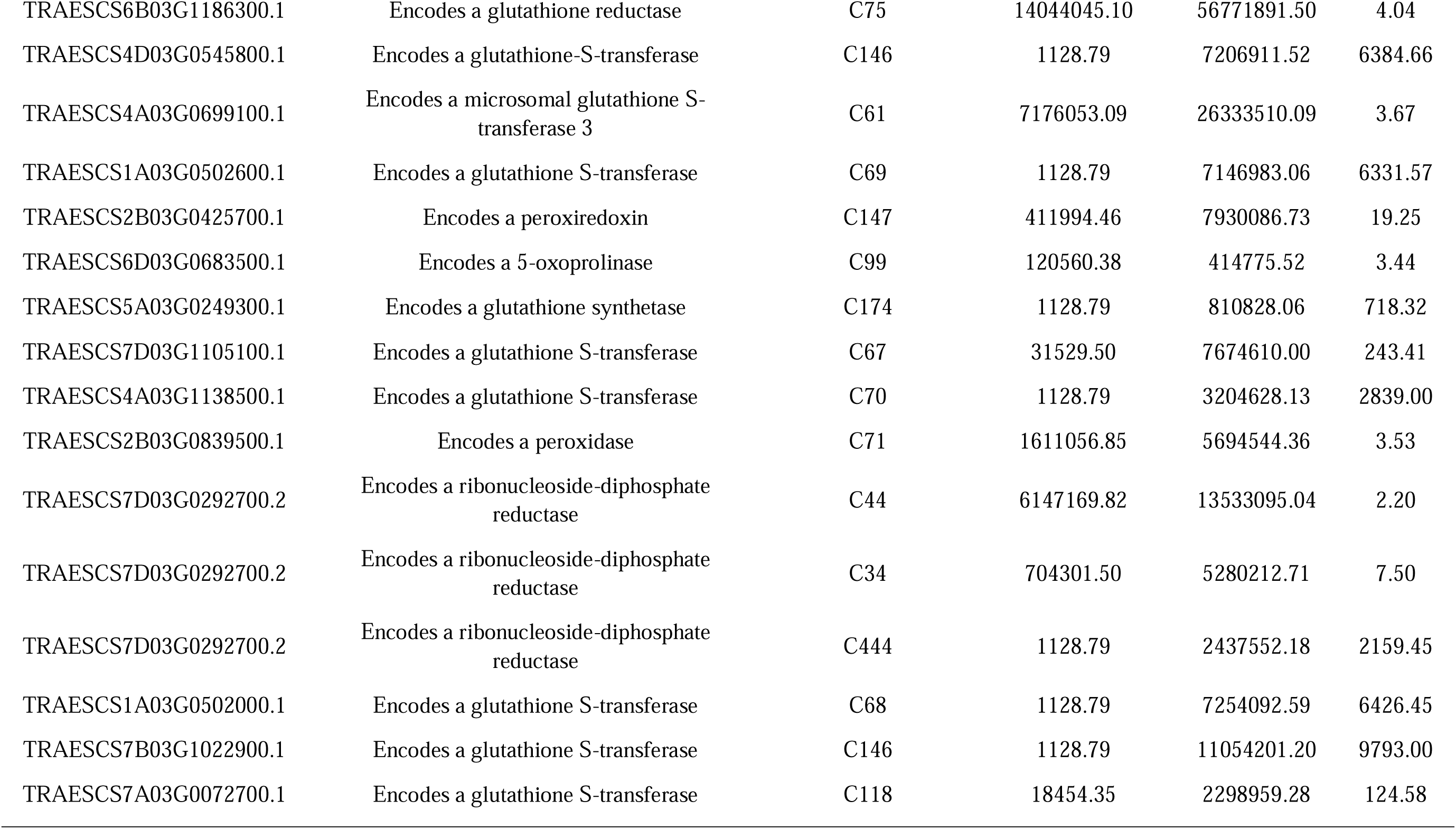
S-nitrosylated proteins involved in GSH biosynthesis and metabolism in *Arabidopsis* and wheat following GSNO treatment.

| Arabidopsis |  |  |  |  |  |  |
|---|---|---|---|---|---|---|
|  |  |  |  | Peptide intensity |  |  |
| Gene ID | Protein name | Gene function | S-nitrosylated site | 0 h | 0.5 h | fold |
|---|---|---|---|---|---|---|
| AT5G02790.1 |  | Encodes glutathione transferase | C37 | 183732.22 | 471579.19 | 2.57 |
| AT2G25080.1 | GPX1 | Encodes glutathione peroxidase | C111 | 7775778.37 | 27010647.40 | 3.47 |
| AT4G35000.1 | APX3 | Encodes a microsomal ascorbate peroxidase APX3 | C31 | 4330819.95 | 3536590.33 | 0.82 |
| AT2G24200.2 | LAP1 | Encodes a leucine aminopeptidase | C305 | 5449640.57 | 7153087.06 | 1.31 |
| AT1G23820.1 | SPDS1 | Spermidine synthase | C346 | 541604.31 | 1670660.34 | 3.08 |
| AT1G59700.1 | GSTU16 | Encodes glutathione transferase | C194 | 2274894.43 | 2756020.84 | 1.21 |
| AT1G70310.1 | SPDS2 | Spermidine synthase. | C326 | 327021.21 | 2206801.45 | 6.75 |
| AT2G31570.1 | GPX2 | glutathione peroxidase GPx | C41 | 54478743.92 | 65739564.92 | 1.21 |
| AT5G16710.1 | DHAR3 | Encodes a dehydroascorbate reductase | C66 | 37945290.90 | 65078882.73 | 1.72 |
| AT5G16710.1 | DHAR3 | Encodes a dehydroascorbate reductase | C69 | 698876.02 | 2778452.03 | 3.98 |
| AT1G19570.1 | DHAR1 | Encodes a dehydroascorbate reductase | C20 | 25653081.36 | 68659378.75 | 2.68 |
| AT4G30920.1 | LAP2 | Encodes LAP2, an aminopeptidase | C209 | 89558.05 | 825826.87 | 9.22 |
| AT5G35790.1 | G6PD1 | Encodes a glucose-6-phosphate dehydrogenase | C149 | 5740703.62 | 10439192.79 | 1.82 |
| AT3G54660.1 | GR2 | Encodes glutathione reductase | C486 | 16823665.53 | 5424961.16 | 0.32 |
| AT2G21790.1 | RNR1 | Encodes large subunit of ribonucleotide reductase | C44 | 1277519.75 | 2050573.22 | 1.61 |
| AT3G24170.3 | GR1 | Encodes a cytosolic glutathione reductase | C73 | 51909407.57 | 59620434.45 | 1.15 |
| AT4G35970.1 | APX5 | Encodes a microsomal ascorbate peroxidase APX5 | C305 | 4330819.95 | 3536590.33 | 0.82 |
| AT1G75270.1 | DHAR2 | One of three functional dehydroascorbate reductases | C20 | 2058594.15 | 5773987.18 | 2.80 |
| wheat |  |  |  |  |  |  |
|  |  |  |  | Peptide intensity |  |  |
| Gene ID |  | Gene function | S-nitrosylated site | 0 h | 1 h | fold |
| TRAESCS4B03G0390700.1 |  | Encodes a glutathione peroxidase | C92 | 1005712.05 | 1184858.71 | 1.18 |
| TRAESCS4D03G0493400.1 |  | Encodes a glutathione-S-transferase | C41 | 1771972.54 | 14212013.15 | 8.02 |
| TRAESCS7D03G0069600.1 |  | Encodes a glutathione S-transferase | C118 | 705235.21 | 4722263.86 | 6.70 |
| TRAESCS4B03G0557600.1 |  | Encodes a glutathione-S-transferase | C38 | 23052.34 | 3071917.39 | 133.26 |
| TRAESCS4B03G0557600.1 |  | Encodes a glutathione-S-transferase | C41 | 23052.34 | 3071917.39 | 133.26 |
| TRAESCS7A03G1156600.1 |  | Encodes a glutathione-S-transferase | C67 | 1128.79 | 6689149.54 | 5925.97 |
| TRAESCS6B03G1186300.1 |  | Encodes a glutathione reductase | C357 | 1128.79 | 2740280.49 | 2427.63 |
| TRAESCS6B03G1186300.1 |  | Encodes a glutathione reductase | C75 | 14044045.10 | 56771891.50 | 4.04 |
| TRAESCS4D03G0545800.1 |  | Encodes a glutathione-S-transferase | C146 | 1128.79 | 7206911.52 | 6384.66 |
| TRAESCS4A03G0699100.1 |  | Encodes a microsomal glutathione S-transferase 3 | C61 | 7176053.09 | 26333510.09 | 3.67 |
| TRAESCS1A03G0502600.1 |  | Encodes a glutathione S-transferase | C69 | 1128.79 | 7146983.06 | 6331.57 |
| TRAESCS2B03G0425700.1 |  | Encodes a peroxiredoxin | C147 | 411994.46 | 7930086.73 | 19.25 |
| TRAESCS6D03G0683500.1 |  | Encodes a 5-oxoprolinase | C99 | 120560.38 | 414775.52 | 3.44 |
| TRAESCS5A03G0249300.1 |  | Encodes a glutathione synthetase | C174 | 1128.79 | 810828.06 | 718.32 |
| TRAESCS7D03G1105100.1 |  | Encodes a glutathione S-transferase | C67 | 31529.50 | 7674610.00 | 243.41 |
| TRAESCS4A03G1138500.1 |  | Encodes a glutathione S-transferase | C70 | 1128.79 | 3204628.13 | 2839.00 |
| TRAESCS2B03G0839500.1 |  | Encodes a peroxidase | C71 | 1611056.85 | 5694544.36 | 3.53 |
| TRAESCS7D03G0292700.2 |  | Encodes a ribonucleoside-diphosphate reductase | C44 | 6147169.82 | 13533095.04 | 2.20 |
| TRAESCS7D03G0292700.2 |  | Encodes a ribonucleoside-diphosphate reductase | C34 | 704301.50 | 5280212.71 | 7.50 |
| TRAESCS7D03G0292700.2 |  | Encodes a ribonucleoside-diphosphate reductase | C444 | 1128.79 | 2437552.18 | 2159.45 |
| TRAESCS1A03G0502000.1 |  | Encodes a glutathione S-transferase | C68 | 1128.79 | 7254092.59 | 6426.45 |
| TRAESCS7B03G1022900.1 |  | Encodes a glutathione S-transferase | C146 | 1128.79 | 11054201.20 | 9793.00 |
| TRAESCS7A03G0072700.1 |  | Encodes a glutathione S-transferase | C118 | 18454.35 | 2298959.28 | 124.58 |

**Supplementary Table S6.** Intensity of S-nitrosylated sites containing Cys 73 of *Arabidopsis* GR1 and Cys 75 of wheat GR1 following GSNO treatment.

| *Arabidopsis* : AT3G24170.3-C73 |  |  |
|  | 0 h | 0.5 h |
|---|---|---|
| Rep.1 | 55620518.00 | 104873280.20 |
| Rep.2 | 38642221.77 | 73986775.47 |
| Rep.3 | 61465482.95 | 1247.69 |
| Average | 51909407.57 | 59620434.45 |
| ±SD | 11855571.19 | 53891832.56 |
| Wheat : TRAESFLD6B01G468300.1-C71 |  |  |
| (Chinese Spring: TRAESCS6B03G1186300.1-C75) |  |  |
|  | 0 h | 1 h |
| Rep.1 | 7048261.12 | 1128.79 |
| Rep.2 | 1128.79 | 112451673.70 |
| Rep.3 | 35082745.38 | 57862872.00 |
| Average | 14044045.10 | 56771891.50 |
| ±SD | 18557631.87 | 56233210.31 |

